# An electro-optical bead-nanochip technology for the ultrasensitive and multi-dimensional detection of small extracellular vesicles and their markers

**DOI:** 10.1101/2022.04.11.487936

**Authors:** Tomás Dias, Ricardo Figueiras, Susana Vagueiro, Renato Domingues, Yu-Hsien Hung, Elnaz Persia, Pierre Arsène

**Affiliations:** Mursla Ltd, Cambridge U.K.

## Abstract

Small extracellular vesicles (sEVs), including exosomes, are enriched in multiomics information mirroring their parental cells. They have been investigated in health and disease and utilised in several applications from drug discovery to diagnostics. In disease diagnostics, sEVs can be sampled via a blood draw, enabling the convenient liquid biopsy of the tissue they originate from. However, few applications with sEVs have been translated into clinical practice.

We developed a Nanoparticle EXOsome Sensing (NEXOS) technology, for the ultrasensitive and multi-dimensional detection of sEVs. NEXOS comprises two methods: a novel nanoelectronics method, E-NEXOS, and a high-throughput optical detection method, O-NEXOS. Both methods share the same steps for the immunocapture and antibody-labelling of sEVs and can be combined to derive differentiated detection parameters.

As a proof of concept, we show the analytical detection and sensitivity of these methods in detecting pre-prepared cancer cell-derived CD9^+^CD81^+^ and CD9^+^HER2^+^ sEVs. Both sEV populations were diluted in PBS and spiked in processed plasma. We also provide a novel approach for the determination of target sEVs (TEVs), target epitopes in sEVs (TEPs), and epitopes per target sEV, as yet unseen from current and emerging technologies.

Further, we demonstrate the higher sensitivity of O-NEXOS compared to the gold standard techniques, as well as demonstrating that E-NEXOS possesses commensurate sensitivity whilst only being powered by 36 nanogap-based sensors per nanochip.

Finally, this manuscript lays the groundwork for a scalable electronics miniaturization of E-NEXOS nanochip with millions of nanogap-based sensors for the translation of NEXOS into standard clinical practice.

## INTRODUCTION

Small extracellular vesicles (sEVs), including exosomes, have emerged as important mediators of intercellular communication [1, 2]. sEVs contribute to a wide range of biological processes in health [3] and disease [4] and are being widely investigated in both therapeutic and diagnostic applications [5, 6].

sEVs are 50 – 150 nm membrane-encapsulated particles that are secreted by virtually all cells, are present in every fluid in the organism and are enriched in proteins, nucleic acids, lipids and metabolites. Since the molecular composition of sEVs mirrors their parental cells [7, 8], they are ideal biomarkers for the development of novel diagnostic tests.

For example, many tumours have been shown to release sEVs into the peripheral blood of corresponding patients long before malignant cells disseminate from the primary tumour [9]. As such, sEVs are increasingly recognized as a promising source of biomarkers to detect many different diseases (particularly cancer and degenerative diseases), accessible from minimally to non-invasive liquid biopsies [2, 10].

While the field is gaining momentum, there is only one commercialised test offering clinical insight: ExoDx Prostate IntelliScore (EPI). This test relies on RNA signatures, typically recovered in sEV preparations derived from urine, and frequently termed as exosomal RNAs [11].

We hypothesise that this is partly because known disease biomarkers carried by sEVs are lacking. Furthermore, detection technologies require higher sensitivity, particularly in the detection of sEV proteins, metabolites and lipids, where no amplification is possible as it is with nucleic acids [5]. Interestingly, a pioneering large-scale discovery analysis of sEV proteomics based on Mass-Spectrometry, demonstrated unique proteomic profiles in blood-derived sEVs across distinct cancer types, validating that unique cancer signatures are in fact contained in sEVs of different origins [12]. The workflow, however, needs further validation and the study does not hint towards targeted detection techniques for translation into clinical assays. Moreover, in a recent study, it was predicted that current bulk sEV detection systems are approximately 10^4^ -fold too insensitive to detect early-stage human cancers of ∼1 cm^3^, whilst emerging sEV methods will allow blood-based detection of cancers of <1 mm^3^ in humans [13].

Currently, commercially available methods for the detection and characterisation of sEVs predominantly consist of variations of Nanoparticle Tracking Analysis (NTA), Tunable Resistive Pulse Sensing (TRPS), Enzyme-Linked Immunosorbent Assays (ELISAs), Western Blot (WB) or bead-based flow cytometry (FCM). These techniques have been adapted from more conventional approaches that are generally too insensitive for early-stage diagnosis of complex diseases. More recently, Single-Particle Interferometric Reflectance Imaging Sensor (SP-IRIS) and Imaging or Nano Flow Cytometry have emerged. However, these techniques are mostly geared towards academic research, are burdened with high equipment and assay costs, and possess at least one of low specificity, low sensitivity, and low throughput capabilities [5, 14].

Several novel academic sEV biosensing approaches have been reported with improved analytical sensitivity over the conventional methods, but these require further validation and scaling analysis for clinical translation [15-19].

In this study, we present a hybrid bead-nanochip technology for the ultrasensitive and multi-dimensional electro-optical detection of sEVs. This technology involves capturing sEVs in a bead-based magnetic immunoassay, isolating the captured sEVs from a biofluid, and sensing the target sEVs via labelling gold nanoparticles (GNPs). We named this novel nanoelectronics method Electro-NEXOS (E-NEXOS). Moreover, we show that the magnetic immunoassay can be combined with the optical reporting of sEVs, named Optical-NEXOS (O-NEXOS), as an alternative to using labelling GNPs.

Both E-NEXOS and O-NEXOS share the same steps for the antibody labelling and immunocapture of sEVs but they differ in the detection method, as implied by their name. Additionally, they offer complementary information to one another which is explored further in this manuscript.

Here, we show the working principle of the NEXOS technology and the parameters we optimised for the development of the methods.

Additionally, using pre-prepared MCF-7 sEVs, we demonstrate the analytical detection of CD9^+^CD81^+^ MCF-7 sEVs with E-NEXOS and O-NEXOS, at different sEV concentrations. Furthermore, we present a workflow for the determination of target sEVs (TEVs) and target epitopes (TEPs) in sEV preparations with E-NEXOS and O-NEXOS methods, respectively. TEVs represent sub-populations of sEVs displaying biomarkers of interest and TEPs are the biomarkers of interest in a given sample containing sEVs.

We then validate the developed methods on sEVs prepared from another cell line, namely BT-474. We show the analytical detection of CD9^+^CD81^+^ and CD9^+^HER2^+^ BT-474 sEVs, discuss the sensitivity of the methods and determine the corresponding concentration of TEVs and TEPs. In addition to that, we show the synergy of the methods in determining, for the first time, the number of target epitope markers per target sEV.

We then compare the sensitivity of E-NEXOS and O-NEXOS with that of established technologies, namely Fluorescence-ELISA, SP-IRIS and Imaging Flow Cytometry (IFC) in the detection of BT-474 sEVs.

Finally, we lay the groundwork for the scalable electronics miniaturisation of E-NEXOS, hinting towards the optimisations that can translate sEV based diagnosticsinto clinical practice.

## RESULTS

### NEXOS working principle

The general workflow of the NEXOS technology is illustrated and summarised in **Fig. 1**. Starting from samples containing sEVs, target sEVs are labelled with “detection” biotinylated antibodies, recognising any target surface protein on sEVs (**Fig. 1A**). Next, using the sandwich principle as in an ELISA, magnetic beads (MBs) pre-coated with “capture” antibodies capture and purify the antibody-labelled target sEVs from primary body fluids, or from cell culture’s conditioned media. Unbound detection antibodies and non-target sEVs are removed upon washing the MB-bound sEVs in a magnetic field (**Fig. 1B**).

**Fig. 1.**
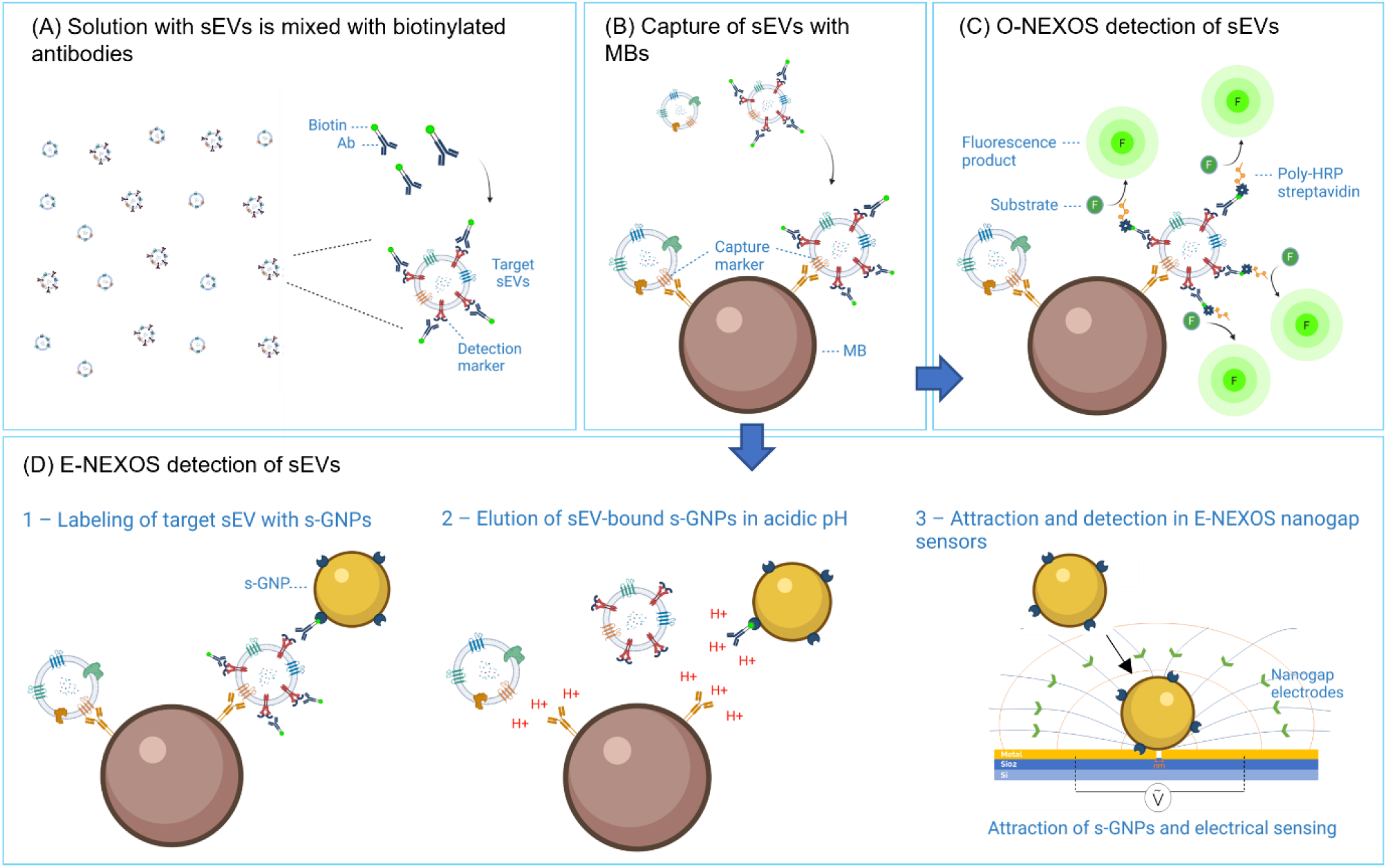
NEXOS workflow for the detection of sEVs. (A) Biotinylated (detection) antibodies recognise and bind to target surface proteins (detection markers) on sEVs. (B) Magnetic Beads (MBs) pre-coated with (capture) antibodies, capture (detection) antibody-labelled target sEVs and non-labelled sEVs. (C) O-NEXOS detection of target sEV epitopes is performed by the stepwise addition of poly-HRP streptavidin, fluorescence substrate and by the measurement of the resultant fluorescence product. (D) Alternatively, 1 – E-NEXOS is performed by adding streptavidin-GNPs (s-GNPs) to the mixture of MB-captured sEVs. sEV-bound GNPs are recovered by washing the MBs-sEVs-s-GNPs complexes in a magnetic field. 2 – s-GNPs are eluted by dissolving the complexes in pH 2.6, then restoring it back to pH 7. 3 – Buffer containing s-GNPs is exchanged by centrifugation to ultra-pure water or 1 % PBS and, finally, the s-GNPs are attracted and detected in an E-NEXOS nanochip.

A polymerised form of horseradish peroxidase (HRP) bound to streptavidin (polyHRP-strep) is incubated with the MBs-captured sEVs and, after washing of unbound polyHRP-strep, a fluorescent substrate is added from which a signal is acquired in O-NEXOS (**Fig. 1C**).

Alternatively, streptavidin-coated GNPs (s-GNPs) of 200 nm are incubated with the MBs-captured sEVs and, upon binding, form a complex comprised of MBs-sEVs-s-GNPs (**Fig. 1D – 1**). Unbound s-GNPs are washed away in a magnetic field, and sEVs-bound s-GNPs are eluted from the MBs (**Fig. 1D – 2**). Finally, the recovered s-GNPs are washed and the buffer is exchanged with either ultra-pure water or 1 % PBS, and the indirect detection of target sEVs is enabled via the attraction and detection of eluted s-GNPs in the E-NEXOS nanogap-based sensors (**Fig. 1D – 3**).

The optical sensing principle of O-NEXOS has been similarly developed by others [20]. Furthermore, in a previous study we demonstrated the capabilities of O-NEXOS in detecting serum and plasma sEVs, alongside reporting it as a magnetic bead ELISA system [21]. For these reasons, we show in this manuscript the novel developments and optimizations of E-NEXOS. Yet, we apply both methods in the detection of sEVs and demonstrate how they can be utilised to derive differentiated measurement parameters.

### sEVs preparation and characterisation

For the proof of concept demonstration of NEXOS, we used sEVs prepared from the conditioned media of MCF-7 (ATCC HTB-22), a low-grade luminal A non-metastatic breast cancer cell line and BT-474 (ATCC HTB-20), an HER-2 overexpressing breast cancer cell line. We used these cell lines since they have been widely utilised in sEV research studies [22-24]. Furthermore, we chose BT-474 as it overexpresses a popular biomarker with implications in cancer, HER2 [25, 26].

In brief, MCF-7 and BT-474 cells were raised in media supplemented with fetal bovine serum (FBS) and, upon reaching 60 – 70 % confluence, cells were washed in PBS and maintained in serum-free media for 48 h. Media were harvested and sEVs prepared by a series of methods combining differential centrifugation, microfiltration, and size exclusion chromatography (SEC). For further information on the preparation of sEVs, please see Materials and Methods. To validate the obtained sEVs for the present study, the sEVs were characterised according to current standards in the field [27]. Particle numbers were quantified by NTA (**Fig. 2A**) and the presence of the sEV specific antigens, CD9, CD81, CD63 and TSG101 was confirmed by WB (**Fig. 2B**). The spherical shape and lipid bilayer membrane was established by Transmission Electron Microscopy (TEM) (**Fig. 2C**) and the presence of CD9, CD81 and CD63 was further confirmed by bead-based Flow Cytometry (FCM) (**Fig. 2D** and **Fig. 2E**). HER2 antigen was not detected in MCF-7 sEVs, while it was confirmed to be present in sEVs isolated from BT-474, again by WB and FCM (**Fig. 2B** and **Fig. 2E**).

**Fig. 2.**
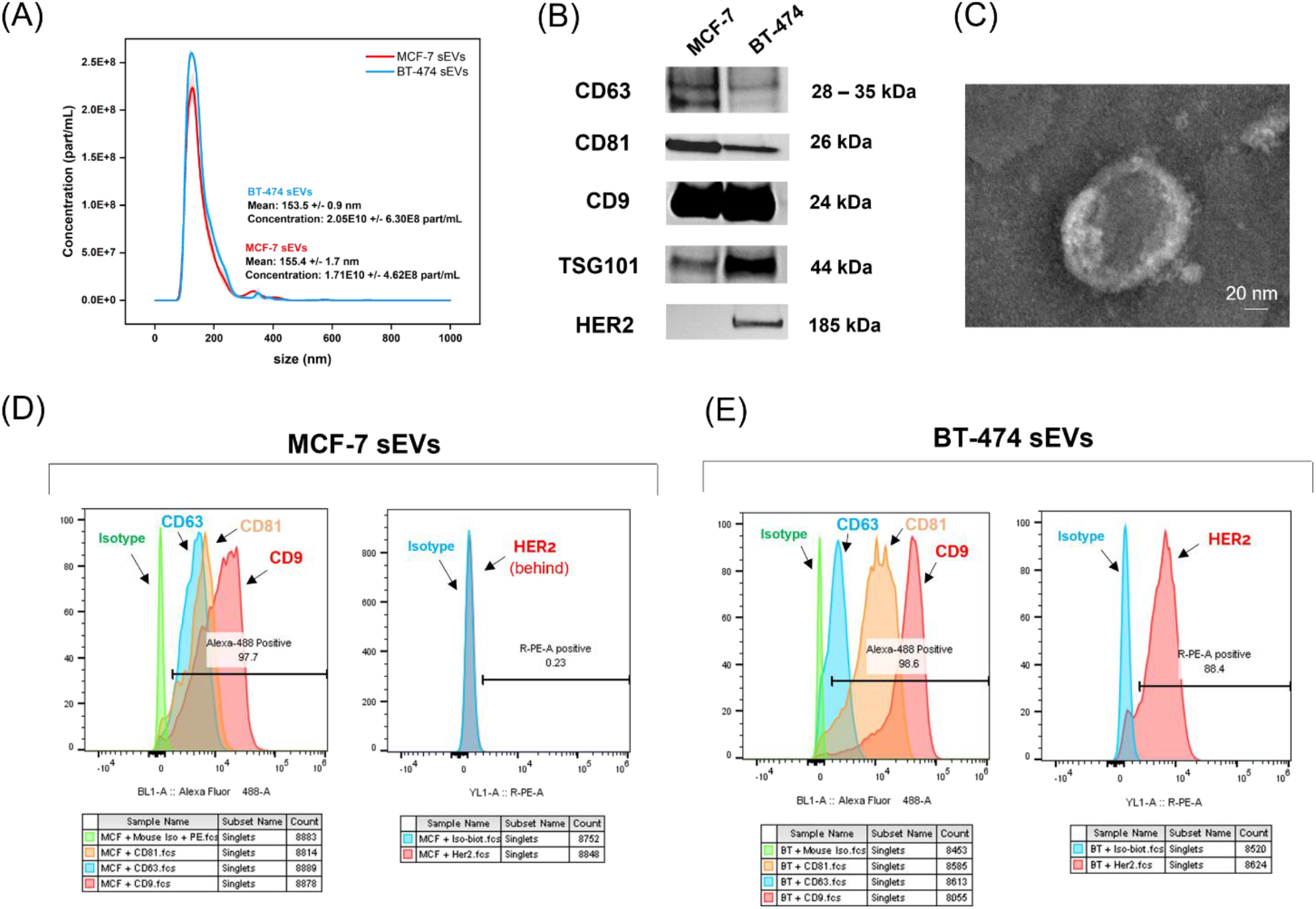
Characterisation of MCF-7 and BT-474 sEVs. (A) Average diameter and particle concentration of pre-prepared MCF-7 and BT-474 sEVs, determined by NTA (n = 3). (B) Western blot analysis of CD63, CD81, CD9, TSG101 and HER2 in lysates of MCF-7 and BT-474 sEV preparations. (C) Size and morphological characterisation of BT-474 sEVs by Transmission Electron Microscopy (TEM). (D-E) Determination of CD63, CD81, CD9 and HER2 proteins in MCF-7 and BT-474 sEVs by Bead-based Flow Cytometry (FCM).

### The design, sensing principle and optimisation of E-NEXOS

In **Fig. 1**, the workflow of NEXOS technology was summarised without clarifying how the labelling s-GNPs are attracted and detected in E-NEXOS nanochip (**Fig. 1D – 3**). This section therefore aims to clarify this. E-NEXOS is a novel method and results from our long-term vision of developing nanochips with more than 1,000,000 nanogap-based sensors for the ultra-sensitive detection of sEVs. A nanogap-based sensor is achieved by separating two nanometer-sized gold electrodes with a nanogap. Each sensor quickly attracts nanoparticles, s-GNPs in this study, in the vicinity via a dielectrophoretic (DEP) force. Nanoparticle sensing is performed on the connected sensors after completing an electrical activation step. Extended explanations of this method are provided in Supplementary Information and in supporting patent references [28, 29].

The current generation of E-NEXOS nanochips resulted from various steps of fabrication and parameter optimisations which we present next. We started by developing a 12 × 12 mm^2^ nanochip matrix consisting of 36 gold sensors. The nanochip fabrication was processed according to the standards in the semiconductor industry (see Materials and Methods).

The sensors are distributed in 4 sensing areas with 9 sensors each (**Fig. 3A**). Each sensor is made of 3 parallel electrode pairs (**Fig. 3B**) as a method to triplicate the attraction surface area. The electrode pairs are made of a layer of Ti5/Au50 (thicknesses in nm), and have a width of 300 nm, a height of 35 nm and a nanogap separation of 50 nm (**Fig. 3C**).

**Fig. 3.**
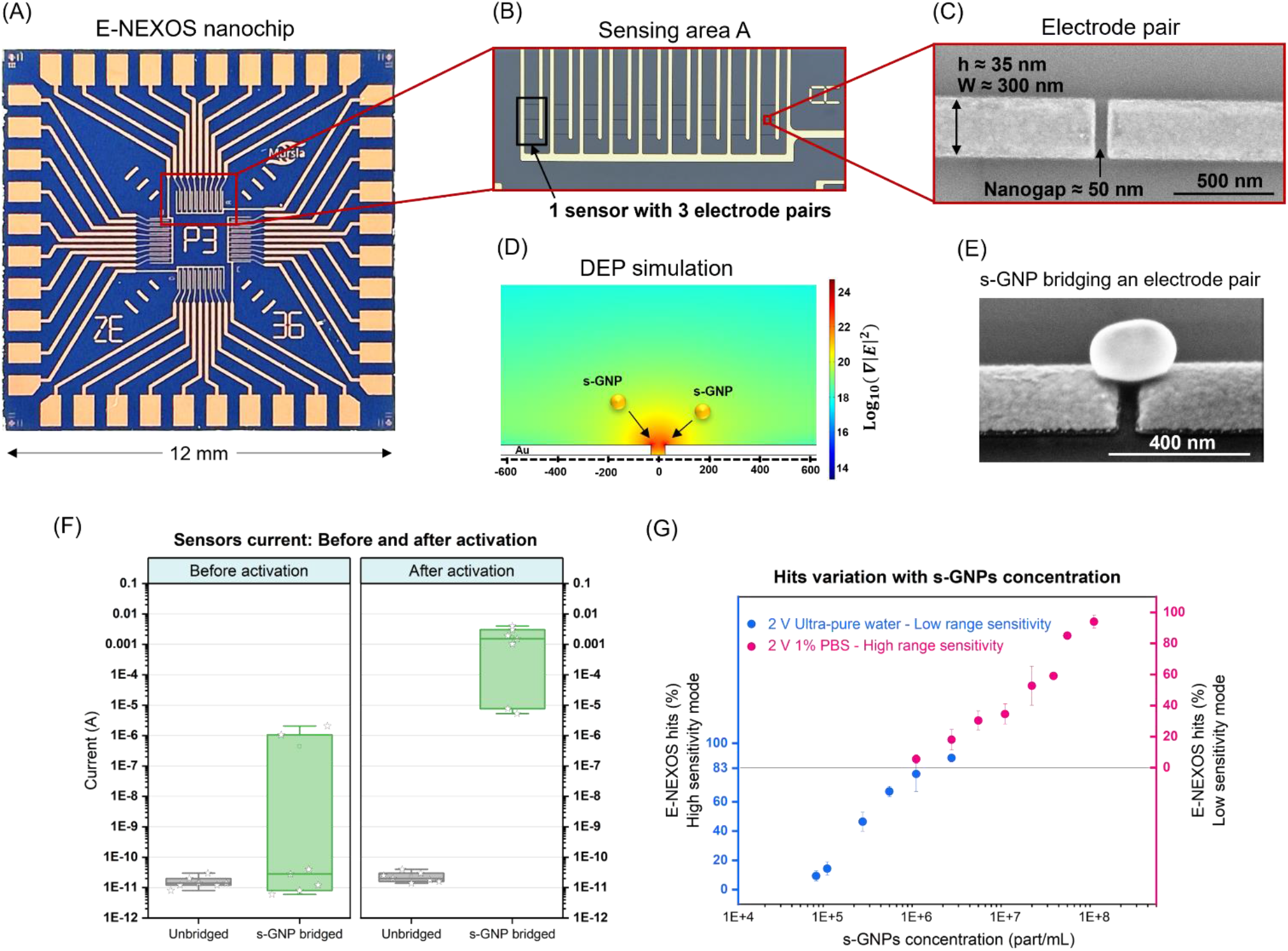
E-NEXOS design and sensing principle. (A) Optical microscopy of an E-NEXOS nanochip with 36 sensors divided into 4 sensing areas. (B) Optical microscopy of a sensing area in E-NEXOS nanochip. Each area contains 9 sensors with 3 nanogaps each. (C) Characterisation of a nanogap by Scanning Electron Microscopy (SEM). The measured dimensions of the nanogap and gold electrodes are described. (D) Simulation of the gradient of the electric field intensity ∇ |*E*^2^| in a nanogap, using the nanogap dimensions and electrical parameters used experimentally. Nanoparticles placed under this field will experience a positive Dielectrophoresis (DEP) force, being attracted to the nanogap. (E) SEM characterisation of an electrode pair separated by a nanogap after attracting streptavidin-GNPs (s-GNPs) and showing an attracted s-GNP bridging the nanogap. (F) Current measured on unbridged nanogaps and on nanogaps after trapping s-GNPs, before and after activation with 5 V DC (n = 7). (G) Determination of the percentage of hits in E-NEXOS in function of the concentration of s-GNPs for E-NEXOS low range sensitivity and high range sensitivity modes (n = 4).

Then, using the geometric parameters described above, we simulated the cross-sectional gradient of the electric field intensity, ∇ |*E*^2^|, generated by 2 V AC at 100 kHz in the vicinity of a nanogap using COMSOL Multiphysics software.

The simulation showed that the highest ∇ |*E*^2^| is at the edge of the electrodes, and it decreases proportionally to the distance from the nanogap (**Fig. 3D**). An s-GNP placed under this electric field experiences a positive DEP force, therefore being attracted to the nanogap (see **Fig. S1** and additional Supplementary Information).

Accordingly, we confirmed by Scanning Electron Microscopy (SEM) that the s-GNPs are trapped in the nanogaps after the attraction step described above. This enabled the connection of the electrode pairs in the nanochip (**Fig. 3E**).

Once connected, the liquid was removed and an activation step at 5 V DC was performed. It fuses the pair of electrodes with the nanoparticles, and short-circuits the electrode pairs with s-GNPs. This promotes a current shift from pA to up to mA, a nine-order increase in signal magnitude and resulted in the unbiased detection of s-GNPs (**Fig. 3F**).

We describe a sensor as ‘connected sensor’ or ‘hit’ when the measured current reached at least 1 µA after both attraction and activation steps. To eliminate any unlikely fabrication variants, it was verified that the current was in the pA range prior to this.

Then, we determined the total percentage (%) hits in a nanochip as in equation 1:

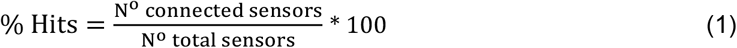

Following the determination of % hits, we investigated different experimental parameters across various s-GNP concentrations. These parameters included the size of s-GNPs, the applied attraction voltage and the permittivity of the s-GNPs detection media (**Fig. S2A-C**).

From our findings, we selected parameters that resulted in the highest sensitivity: an attraction voltage of 2 V AC at 100 kHz, s-GNPs with a diameter of 200 nm and the elution of s-GNPs in ultra-pure water media. To increase the concentration range of detection, we also considered eluting the s-GNPs in 1 % PBS media, which resulted in two modes of detection: a low range sensitivity and high range sensitivity mode (**Fig. S2-D** and **S2-E**).

Lastly, after optimising the sensing parameters, we aimed to confirm our theoretical scaling assumption that the sensitivity increases with an increasing number of electrode pairs per nanochip. Therefore, we compared the sensitivity of nanochips containing 36 sensors (with 3 electrode pairs per sensor) against a previous generation of nanochips containing only 19 sensors (with 1 electrode pair per sensor). We observed that with an approximately 6x increase in sensor area (108 relative to 19 electrode pairs), the sensitivity achieved was 7.5E4 particles/mL (part/mL). This was 6x higher than with 19 sensors, where the sensitivity was limited to 4.5E5 part/mL (**Fig. S2-F**). It confirmed our scaling assumption that increasing the number of sensors leads to a proportional increase in sensitivity.

The optimisations described in this section enabled the current E-NEXOS nanochip to have a detection range of 4 orders of magnitude and to establish our calibration curve for the determination of s-GNPs bound to sEVs (**Fig. 3G**). This is demonstrated in the following section.

### The integration of sEVs capture with magnetic beads enable the detection of sEVs in E-NEXOS and O-NEXOS

Having developed and optimised the sensing parameters, we then established a method for the capture of sEVs. This linked the recovery of s-GNPs (E-NEXOS) and poly-HRP strep (O-NEXOS) with the sEV concentrations of interest. For this, 2.8 µm MBs were pre-coated with sEVs capturing antibodies, anti-CD9 or anti-CD81.

We determined the antibody coating efficiency on the MBs with a Bicinchoninic acid (BCA) protein assay and observed a good correlation with the coating values estimated by the supplier (**Fig. S3-A**). We estimated the number of antibodies coupled per MB in our reactions to be on average 3.1E5 anti-CD81 antibodies and to 4.5E5 anti-CD9 antibodies per MB (**Table S1**), suggesting that the MBs possessed a high binding affinity for CD81^+^ and CD9^+^ sEVs.

As the antibody coating efficiency was higher for MBs pre-coated with anti-CD9 antibody, we determined the capture rate of target CD9^+^ MCF-7 sEVs at different timepoints in comparison to isotype IgG pre-coated control MBs. We concluded that the best signal-to-noise ratio is achieved at 3h incubation (**Fig. S3-B**).

We selected 3 h as the incubation period in our assays and calculated the capture rate efficiency of the method when capturing a population of enriched CD9^+^ sEVs. We obtained a capture rate of 97 % and concluded that our method showed high efficiency in capturing target sEVs (**Fig. S3-C**). Next, we integrated our bead-capture method with E-NEXOS and O-NEXOS, in line with the illustration presented in **Fig. 1**.

First, sEVs prepared from an MCF-7 culture were serially diluted to samples starting at a concentration of 5E8 part/mL down to 5E6 part/mL. Next, the sEVs were stained with biotinylated anti-CD81 antibody and captured with anti-CD9 MBs or isotype IgG MBs.

In E-NEXOS (simplified illustration in **Fig. 4A**), the captured sEVs were counterstained with 200 nm s-GNPs. After 30 min of incubation, unattached s-GNPs were removed, and the complexes formed by MBs-sEVs-s-GNPs were dissolved in pH 2.6. The s-GNPs were recovered and the buffer containing s-GNPs was substituted to ultra-pure water (low range sensitivity) or 1% PBS (high range sensitivity) for attraction. A A DEP potential of 2 V AC (100 KHz) was used, 1 minute per sensor, in E-NEXOS.

**Fig. 4.**
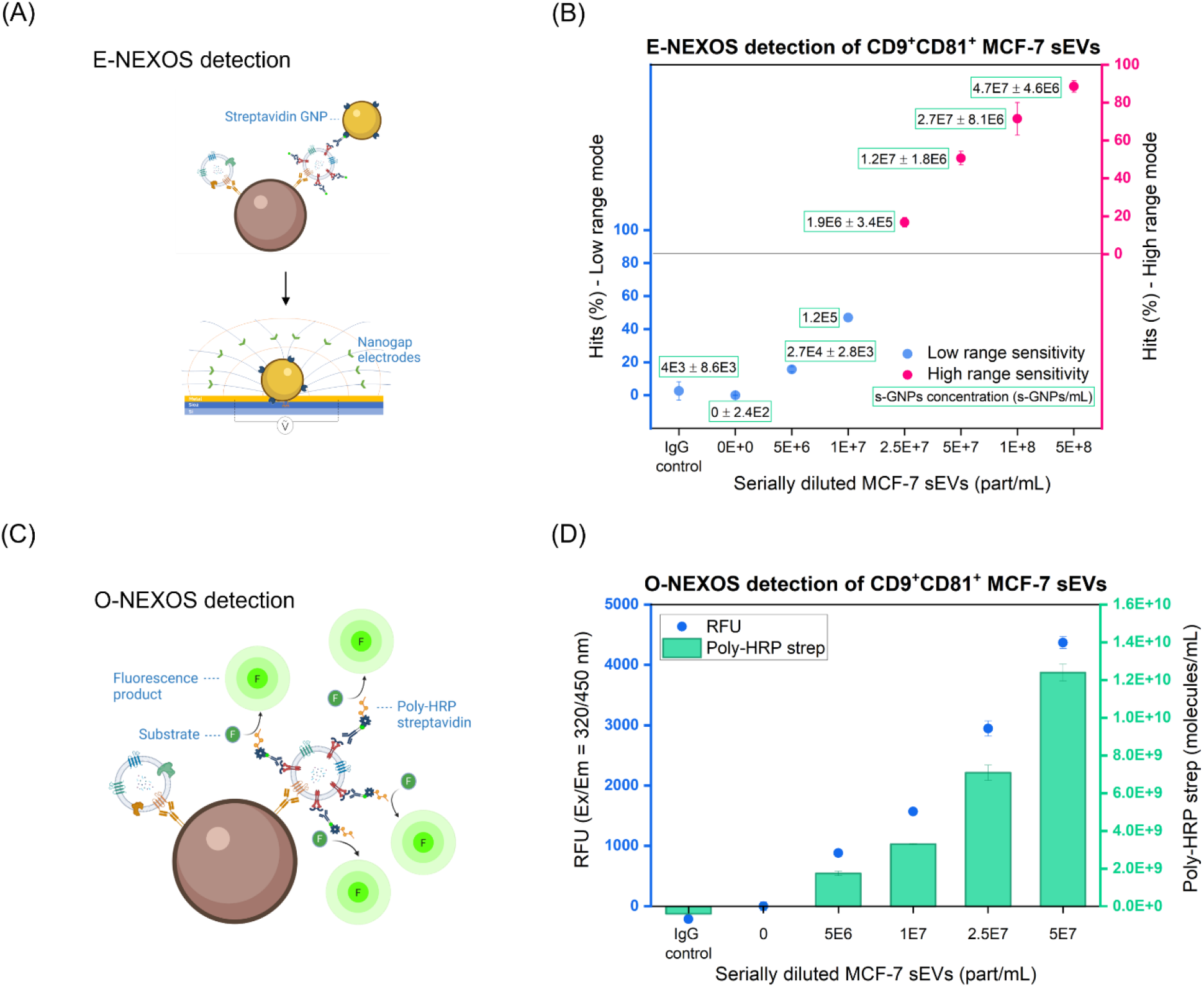
The integration of sEVs capture with magnetic beads enables the detection of sEVs in E-NEXOS and O-NEXOS. (A) Schematic showing sEVs captured on a Magnetic Bead (MB) and integration in E-NEXOS by the labelling of target sEVs with s-GNPs and indirect determination of target sEVs concentrations via an E-NEXOS nanochip. (B) E-NEXOS detection of pre-prepared bulk MCF-7 sEVs by the capture of MCF-7 sEVs with MBs pre-coated with anti-CD9 antibody and detection with s-GNPs targeting CD81 antigens on the sEVs at serially diluted sEV concentrations. The measurements include both low range and high range sensitivity modes of E-NEXOS (n = 4). The measurements included controls with anti-CD9 MBs and without sEVs (blank) and MBs pre-coated with igG antibodies and sEVs at a concentration of 5E8 part/mL (IgG control). (C) Schematic showing sEVs captured on a Magnetic Bead (MB) and integration in O-NEXOS by the labelling of target sEVs with Poly-HRP streptavidin, reaction of Poly-HRP streptavidin with fluorescence substrate and measurement of the resultant fluorescence. (D) O-NEXOS detection of pre-prepared bulk MCF-7 sEVs by the capture of MCF-7 sEVs with MBs pre-coated with anti-CD9 antibody and detection Poly-HRP streptavidin targeting CD81 antigens on the sEVs at serially diluted sEVs concentrations. The graph includes the fluorescence signal detected (left y axis) and the determined number of poly-HRP streptavidin per reaction (right y axis) (n = 4). The measurements included controls with anti-CD9 MBs and without sEVs (blank) and MBs pre-coated with igG antibodies and sEVs at a concentration of 5E8 part/mL (IgG control).

After attraction, the liquid containing s-GNPs was removed and activation of the contacts was completed at 5 V DC for 20 seconds. Finally, sensing was performed with a sweeping voltage from 0 to 5 V DC.

We observed that the number of hits (%) proportionally increases with the concentration of sEVs in both modes of detection (**Fig. 4B**). Using the calibration curve of **Fig. 3G**, the concentration of s-GNPs bound to CD9^+^CD81^+^ sEVs was determined (graph inset values, **Fig. 4B**).

With O-NEXOS (simplified illustration in **Fig. 4C**), instead of incubating the captured sEVs with s-GNPs, we utilized poly-HRP strep for the optical reporting of sEVs. Poly-HRP strep binds to available biotinylated antibodies on the labelled sEVs. After a washing step, a fluorogenic substrate was added and the resulting fluorescent product was quantitated by fluorometry after another washing step. As with s-GNPs, we observed that the fluorescence signal proportionally increased with the concentration of MCF-7 sEVs (left y axis in **Fig. 4D**). Also, with a calibration curve of the generated fluorescence signal in function of the concentration of poly-HRP strep (**Fig. S4**), the concentration of poly-HRP strep bound to sEVs, in O-NEXOS, was estimated (right y axis in **Fig. 4D**).

As observed in **Fig. 4B** and **Fig. 4D**, both E-NEXOS and O-NEXOS exhibit a linear detection response in function of the concentration of target sEVs. Although as of yet we have not tested the sensitivity of both methods, we observed that E-NEXOS and O-NEXOS detect at least CD9^+^CD81^+^ sEVs in a mixed population of sEVs diluted down to 5E6 part/mL.

### Determination of the concentration of target sEVs (TEVs) and target epitopes (TEPs) in samples containing sEVs

Having established the analytical detection of sEVs, we aimed to use E-NEXOS and O-NEXOS to determine the concentration of target sEVs (TEVs) and target biomarker epitopes (TEPs) in samples containing sEVs.

As for E-NEXOS, we postulated that the concentration of TEVs could be determined in if we ascertained how many s-GNPs bind, on average, per target sEV. We hypothesised that the number of s-GNPs that can bind per MB-captured sEV was limited in number, regardless of the number of target epitopes displayed in the target sEV. This hypothesis was formulated due to the relatively large size of s-GNPs (∼ 200 nm), being twice the average size of sEVs (∼ 100 nm). In fact, we determined theoretically that no more than 2 s-GNPs can be bound, at any moment, to an MB-captured sEV (**Table S2**).

Next, we aimed to determine experimentally the number of s-GNPs bound per target CD81^+^CD9^+^ sEV, at different sEV concentrations. In this work, we rely on techniques such as NTA to determine concentration references in sEV preparations. Yet, because sEV preparations such as the ones used in **Fig. 4B** and **Fig. 4D** are heterogenous (containing different sEV populations), these techniques cannot determine the concentration of target sEV populations in these preparations. Therefore, we aimed to enrich for a subpopulation of CD9^+^CD81^+^ sEVs and measure their concentration with NTA for the determination of the number of s-GNPs bound per target CD81^+^CD9^+^ sEV.

Thus, we serially captured and eluted MCF-7 sEVs, first with MBs pre-coated with antibody anti-CD9 and after with MBs pre-coated with antibody anti-CD81, to enrich for a sub-population only comprising (target) CD9^+^CD81^+^ sEVs. As with E-NEXOS, the elution of sEVs between each step was performed in 0.2 M Glycine pH 2.6, following previously described and validated methodologies [30, 31]. Then, the enriched CD9^+^CD81^+^ sEV preparations were serially diluted into samples starting at a concentration of 1E8 part/mL down to 1E6 part/mL and E-NEXOS was used in low range sensitivity and high range sensitivity modes. As observed previously, the percentage of hits (%) proportionally increased with the concentration of sEVs in both modes of detection. We show the concentration of s-GNPs bound per sEV for the different sEV concentrations in **Fig. 5A**. Here, the number of hits is shown as inset values to demonstrate how equivalent data can be displayed in different formats (**graph inset values, Fig. 5A**).

**Fig. 5.**
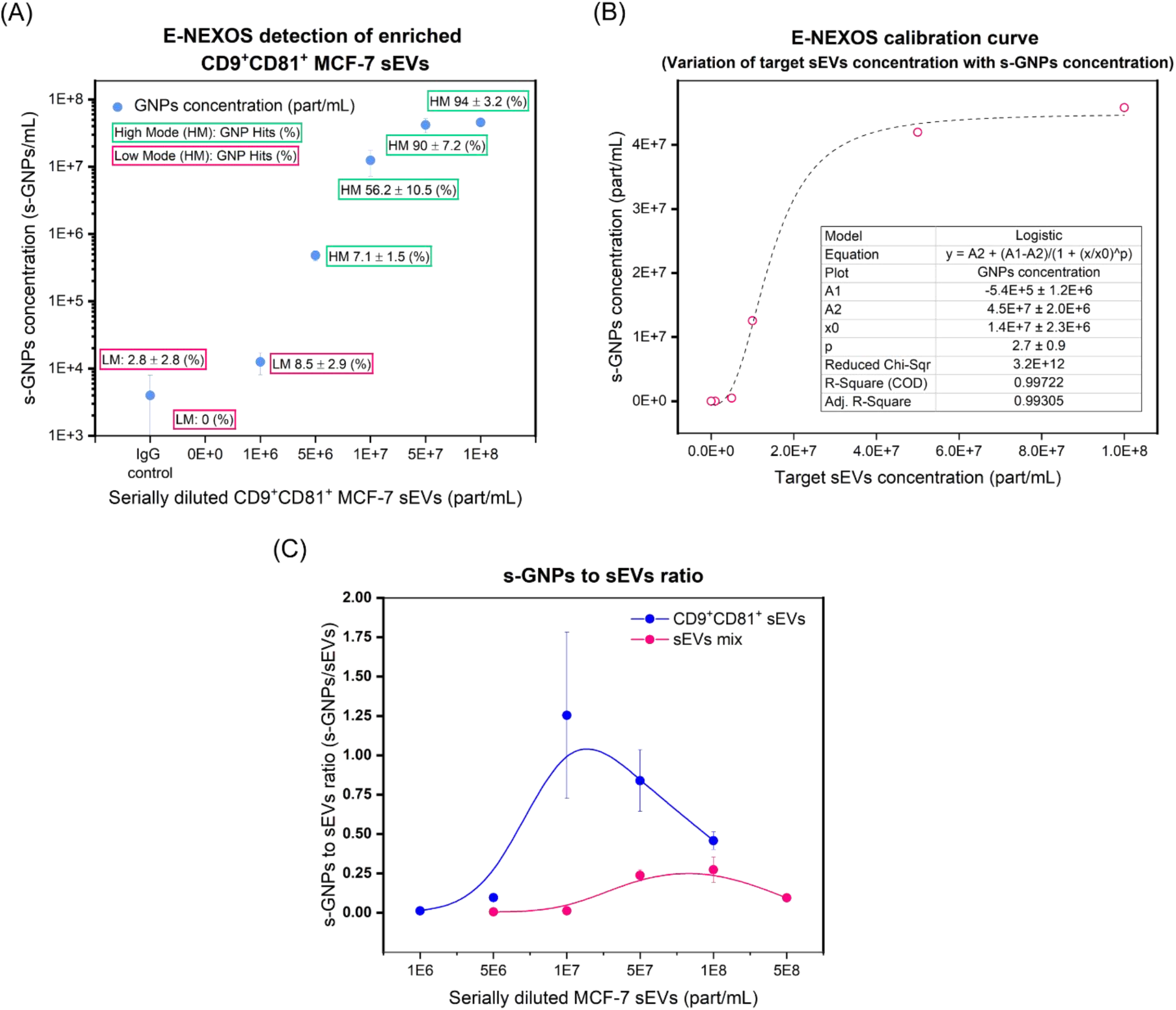
Determination of TEVs. (A) E-NEXOS detection of enriched CD9^+^CD81^+^ MCF-7 sEVs by the capture of CD9^+^CD81^+^ MCF-7 sEVs with MBs pre-coated with anti-CD9 antibody and detection with s-GNPs targeting CD81 antigens on the sEVs at serially diluted sEV concentrations. The measurements include both low range and high range sensitivity modes of E-NEXOS (n = 4). The measurements included controls with anti-CD9 MBs and without sEVs (blank) and MBs pre-coated with igG antibodies and sEVs at a concentration of 5E8 part/mL (IgG control). (B) E-NEXOS was calibrated by pre-enriching CD9^+^CD81^+^ MCF-7 sEVs and detected by E-NEXOS at serially diluted sEV concentrations. With the calibration method described in Fig. 3G, the percentage of hits (%) was translated to the corresponding number of s-GNPs, in this experiment, for the different concentrations of CD9^+^CD81^+^ sEVs measured. A calibration curve enabling the derivation of the concentration of target sEVs (TEVs) was determined in function of the concentration of s-GNPs (n = 4). (C) Determination of the ratio and standard deviation of s-GNPs bound per sEV in a mix of sEVs populations (graph – pink line) and in a population of enriched CD9^+^CD81^+^ sEVs (graph – blue line).

A calibration curve was derived, demonstrating the correlation of the concentration of s-GNPs in an assay with the concentration of target sEVs (TEVs) (**Fig. 5B**).

When comparing the detection of enriched CD9^+^CD81^+^ MCF-7 sEVs (**Fig. 5A**) with the detection performed in **Fig. 4B**, we observed different ratios of s-GNPs bound per sEV (**Fig. 5C**). Additionally, both follow a similar distribution where the ratio of s-GNPs per target sEV increases with increasing sEV concentrations before beginning to decrease. This is possibly due to the amount of sEVs in excess of the fixed concentration of s-GNPs wherein the ratio of s-GNPs bound to sEVs is expected to decrease.

We noted that with sEV concentrations smaller than 7.5E6 part/mL, the ratio of s-GNPs detected per sEV is likely underrepresented due to the current limitations of E-NEXOS nanochip with 36 sensors, in accurately determining low concentrations of s-GNPs.

Furthermore, the maximum ratio of s-GNPs bound per sEV reaches close to 2 s-GNPs per sEV, which is in line with our theoretical prediction (See **Table S2** in supplementary information).

Regarding CD9^+^CD81^+^ sEVs, we achieved a mean ratio of 0.53 ± 0.46 s-GNPs bound per sEV across the different concentrations. However, to better determine the concentration of target sEVs from the concentration of bound s-GNPs, we fit a logistic equation that best describes the relationship between the s-GNPs and target sEVs (TEVs) at different concentrations (**Fig. 5B**). This was subsequently used to determine TEVs in sEV populations obtained from a different cell line.

As for O-NEXOS, we postulated that the concentration of TEPs could be determined by calculating the concentration of poly-HRP strep molecules bound per target epitopes in our reactions. Considering the substantially smaller size of poly-HRP strep molecules (∼ 9.4 nm) when compared with the s-GNPs (200 nm), we estimated that several poly-HRP strep molecules are hypothetically capable of binding to a target sEV, at least one per target epitope counterstained with a biotinylated detection antibody.

Therefore, we hypothesised that O-NEXOS was a suitable indicator for determining the number of TEPs in sEVs, given that such information is complementary to determining TEVs with E-NEXOS. Additionally, given that igG antibodies carry two identical antigen-binding sites, we assumed that each detection antibody could bind to either one or two target epitopes on sEV surface proteins. In turn, poly-HRP strep (at a stoichiometry of 6:1 HRPs per Streptavidin, provided by the supplier), displays up to four biotin-binding sites, therefore binding up to four biotin-antibodies. Thus, in our assay, we assumed a Gaussian distribution for the binding of poly-HRP strep to TEPs as 4.5 ± 2.3 and obtained the equation 2 for the calculation of TEPs:

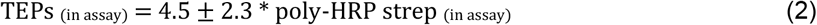

### The determination of TEVs and TEPs can be applied to different sEV populations and their combination reveals a new dimension even in complex samples

Previously, we demonstrated the analytical detection of sEVs alongside strategies to determine the concentration of TEVs and TEPs in mixed sEV preparations, using E-NEXOS and O-NEXOS, respectively.

Aiming to stress the sensitivity of the methods and to validate the methodology to determine TEVs and TEPs in other sEVs preparations, we utilised sEVs isolated from another cell line, BT-474. We targeted a subpopulation of CD9^+^CD81^+^ sEVs, identical to MCF-7 sEVs, but also added a subpopulation of sEVs not present in MCF-7, CD9^+^HER2^+^ sEVs.

This time, the prepared sEVs were serially diluted into different concentrations in PBS, as well as being spiked into processed human plasma. sEVs were stained first with biotinylated antibodies anti-CD81 and anti-HER2 and captured with MBs anti-CD9.

We obtained an analytical detection range in function of the concentration of sEVs (**Fig. S5-A**).

Additionally, we observed a limit of detection (LOD) of 5E6 part/mL and 7.5E5 part/mL for CD9^+^CD81^+^ TEVs and TEPs detection, with a limit of quantification (LOQ) down to 2.5E7 part/mL for TEVs and 7.5E5 part/mL for TEPs. At lower concentrations, the signals overlapped with that of buffer controls (with no sEVs) or after sEV capture with MBs pre-coated with isotype IgG (**Fig. 6A**).

**Fig. 6.**
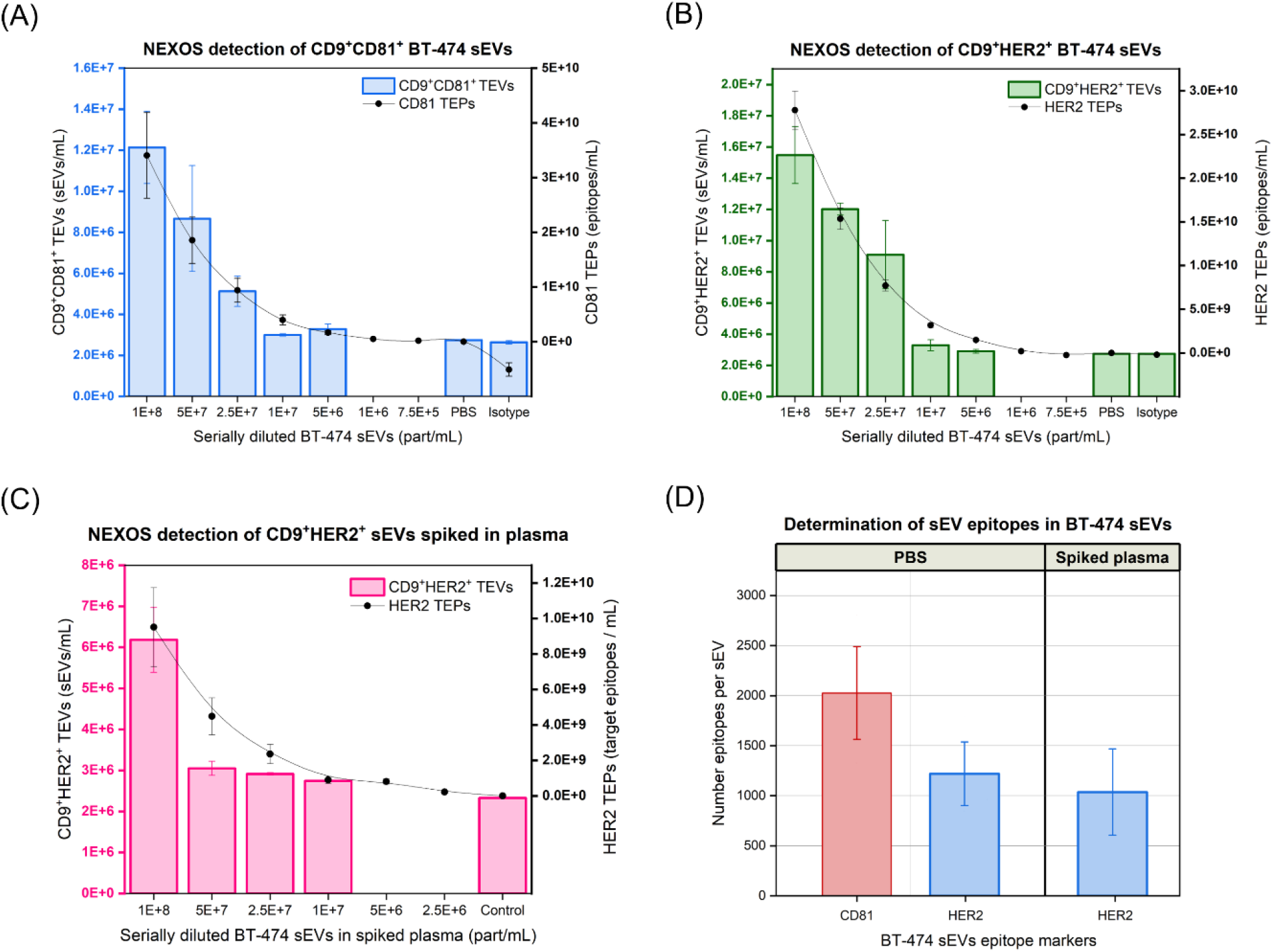
Analytical detection and multi-dimensional characterisation of target sEVs (TEVs), target epitopes (TEPs) and epitopes per sEV in BT-474 sEVs. (A) Analytical detection of CD9^+^CD81^+^ BT-474 sEVs and determination of CD9^+^HER2^+^ TEVs (E-NEXOS) and HER2 TEPs (O-NEXOS) at serially diluted BT-474 sEV concentrations (n = 4). (B) Analytical detection of CD9^+^HER2^+^ BT-474 sEVs and determination of CD9^+^HER2^+^ TEVs (E-NEXOS) and HER2 TEPs (O-NEXOS) at serially diluted BT-474 sEV concentrations (n = 4). (C) Analytical detection of CD9^+^HER2^+^ BT-474 sEVs spiked in processed plasma and determination of CD9^+^HER2^+^ TEVs (E-NEXOS) and HER2^+^ TEPs (O-NEXOS) at serially diluted BT-474 sEV concentrations in the spiked plasma (n = 4). (D) Determination of CD81 and HER2 epitopes per BT-474 sEV, determined from the measurements presented in Fig. 5A, 5B and 5C (n = 4). The measurements included controls with anti-CD9 MBs and without sEVs (PBS) and MBs pre-coated with igG antibodies and sEVs at a concentration of 5E8 part/mL (IgG isotype control).

For HER2 TEVs, we also observed a signal varying in function of the concentration of sEVs (**Fig. S5-B**). The LOD for CD9^+^HER2^+^ TEVs was reached at 5E6 part/mL, and for CD9^+^HER2^+^ TEPs, the LOD was reached at 1E6 part/mL, with an LOQ of 5E6 part/mL TEVs and 1E6 part/mL TEPs (**Fig. 6B**). Notably, higher resolution for both TEVs and TEPs was obtained for the detection with HER2 antibody when compared to CD81 antibody due to the higher unspecific signal in the CD81 controls. Serially diluted sEV preparations spiked in processed plasma were stained with biotinylated anti-HER2 antibody and processed, as previously, for HER2 TEPs and TEVs detection. As before, an analytical relationship between the concentration of sEVs and the signal generated by E-NEXOS and O-NEXOS was observed (**Fig. S5-C**).

For CD9^+^HER2^+^ TEVs, an LOD and LOQ were reached with E-NEXOS system at 1E7 part/mL and for CD9^+^HER2^+^ TEPs, an LOD and LOQ of 2.5E6 part/mL were achieved with O-NEXOS nanochip (**Fig. 6C**). Notably, we observed that the sensitivity for sEVs detection decreases when the sEVs are spiked in processed plasma when compared to the detection of HER2 sEVs in **Fig. 6B**. Nevertheless, this is anticipated given the added complexity of human plasma.

Finally, considering the synergy in determining TEVs and TEPs, we aimed to calculate the number of CD81 and HER2 epitopes per sEV across the three experiments. First, we determined the ratio of TEPs per TEV in the different measurements across the three experiments. Then, we calculated the average of the ratios at sEV concentrations ranging from 1E8 down to 1E7 part/mL. We chose these concentrations as they can be compared across the different experiments.

For BT-474 sEVs diluted in PBS (as shown in **Fig. 6A** and **Fig. 6B**), we determined that the sEVs contained, on average, 2027 ± 535 CD81 epitopes and 1220 ± 368 HER2 epitopes. When BT-474 sEVs were spiked in processed plasma (as shown in **Fig. 6C**), we determined that the spiked sEVs contained on average 1037 ± 522 HER2 epitopes. Notably, we calculated an equivalent number of HER2 epitopes irrespective of the matrix where BT-474 sEVs were eluted in, either PBS or processed plasma. This suggests that the methodology can be applied equally to sEVs in complex samples. The determination of CD81 epitopes on sEVs spiked in processed plasma was not performed because we would also detect the signal originated from sEVs naturally occurring in human plasma and this would not be useful to validate the method.

### Compared to other technologies, O-NEXOS shows the highest sensitivity and E-NEXOS is on-par with the gold standard techniques

We compared the analytical range of NEXOS technology with gold-standard techniques used for the characterization of sEVs, namely Fluorescence-ELISA, IFC and SP-IRIS.

BT-474 sEVs, all from the same batch, were serially diluted to concentrations of 1E10, 5E9, 1E9, 5E8, 1E8, 5E7, 1E7 and 5E6, and compared with the linear sensitivity of NEXOS in detecting BT-474 sEVs.

Like NEXOS, the detection with Fluorescence-ELISA or SP-IRIS utilised capture and detection steps. We chose a commercially available ELISA kit which captures sEVs on the wells of a microtiter plate with high affinity and detects sEVs with anti-CD81 antibody, the ExoELISA-ULTRA CD81 kit. Additionally, this kit was chosen for the comparison as it claimed to be ultra-sensitive.

SP-IRIS was performed by capturing sEVs on ExoView^®^ chips with antibody anti-CD9 and detection with antibodies anti-CD81 and anti-HER2, as in NEXOS.

The exception was Imaging Flow Cytometry (IFC) performed on an ImageStream^®^ X Mk II, where the sEVs were directly labelled with detection antibodies, anti-CD81 and anti-HER2 (without pre-capture with anti-CD9).

We determined the calibration curve of ExoELISA-ULTRA CD81 kit using the standards and instructions provided by the supplier (**Fig. S6-A**). Using the serially diluted BT-474 sEVs, ExoELISA-ULTRA CD81 kit detected sEVs down to a LOD of 5E9 part/mL with an LOQ of 1E10 part/mL (**Fig. S6-B**).

SP-IRIS detected sEVs down to a concentration of 1E7 part/mL, the smallest concentration we tested (**Fig. S6-C** and **Fig. S6-D**). We estimated the LOQs of SP-IRIS as they were not reached at 1E7 part/mL. The determined LOQs were respectively, 9.4E6 part/mL for CD9^+^HER2^+^ sEVs and 4.7E6 part/mL for CD9^+^CD81^+^ sEVs (**Fig. S6-E**).

IFC detected CD81 and HER2 down to an LOQ of 5E7 part/mL for both markers, albeit detecting more signal overall for HER2 (**Fig. S6-F**).

Finally, we compared the LOQ of E-NEXOS and O-NEXOS with the tested techniques.

Overall, O-NEXOS was the most sensitive technique, reaching down to an LOQ of 7.5E5 part/mL, while E-NEXOS already shows a sensitivity comparable to SP-IRIS but superior to IFC or Fluorescent ELISA (**Fig. 7**).

**Fig. 7.**
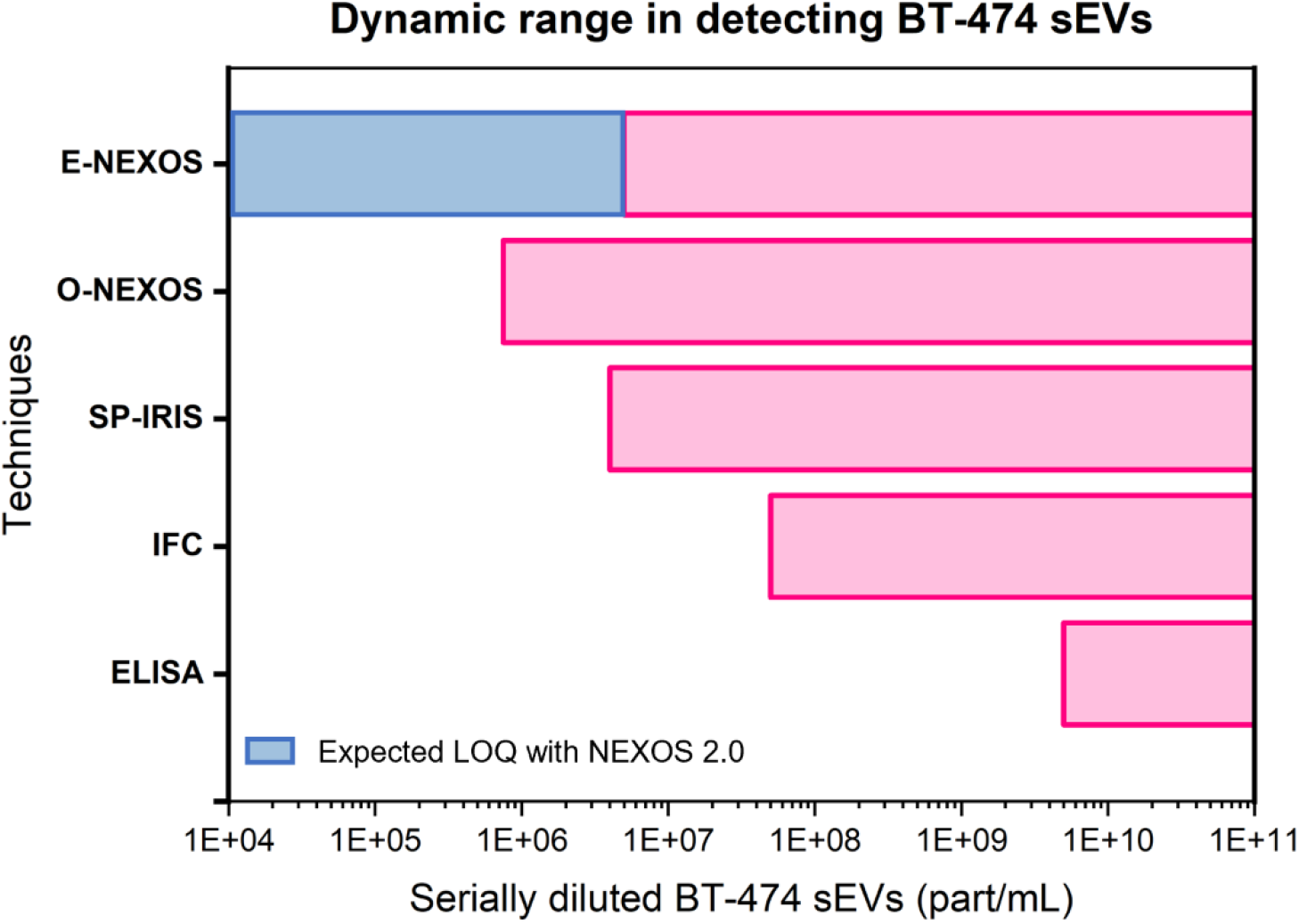
Comparison of different technologies in detecting serially diluted concentrations of BT-474 sEVs. The bars in pink represent the conditions where the highest sensitivity was achieved with each of the tested techniques, namely CD9^+^HER2^+^ sEVs (E-NEXOS) and CD9^+^CD81^+^ sEVs (O-NEXOS). SP-IRIS measured CD9^+^CD81^+^ and CD9^+^HER2^+^ sEVs equally down to 1E7 part/mL, IFC reached the limit of quantification (LOQ) at 5E7 part/mL for either CD81^+^ and HER2^+^ sEVs. ELISA was performed with an ultrasensitive kit reaching down to a LOQ of 5E9 part/mL. The LOQ for SP-IRIS was determined to reach down to 4.7E6 part/mL CD9^+^CD81^+^ sEVs (bar in red). Finally, we show that E-NEXOS can be vastly optimised and reach considerably lower LOQs when compared to the other techniques.

## DISCUSSION

In this study, we present a novel bead-nanochip hybrid technology named NEXOS, which includes two methods: E-NEXOS and O-NEXOS.

Both methods share common steps, however the detection in E-NEXOS is performed on a novel nanoelectronics format whilst O-NEXOS is based on optical detection.

E-NEXOS and O-NEXOS are both complemented by an initial step of sEVs capture on MBs pre-coated with capture antibodies. This step is crucial as it enables selection of only the s-GNPs or poly-HRP strep that are bound to sEVs before detecting the signal generated in E-NEXOS or O-NEXOS. First, we presented the working principle of the technologies (**Fig. 1**), the development of E-NEXOS (**Fig. 3**), and the various parameters we optimised for the detection of sEVs.

Then, we demonstrated the linear detection of the systems in detecting CD9^+^CD81^+^ MCF-7 sEVs (**Fig. 4B** and **Fig. 4D**).

Next, we explored the unique individualities of the systems in determining TEVs and TEPs, respectively in CD9^+^CD81^+^ MCF-7 sEVs (**Fig. 5**). Then, as proof of concept, we validated the determination of TEVs and TEPs on sEVs derived from another cell line, BT-474, by the targeting of CD9^+^CD81^+^ and CD9^+^HER2^+^ sEV populations diluted in PBS or spiked in processed plasma (**Fig. 6A-C**).

Then, we developed a strategy where the two techniques can be combined to derive the number of target epitopes per target sEV. This approach can also be applied if the target sEVs are mixed in heterogeneous sEV samples or eluted in different fluids as demonstrated by the detection of HER2 epitopes per sEV in **Fig. 6D**.

Finally, we compared the sensitivity of the technologies with gold standard techniques (**Fig. 7**).

In NEXOS, the steps involving the labeling of sEVs with detection antibodies, capture of sEVs with MBs, and counterstaining with s-GNPs or poly-HRP strep, are high-throughput, occurring in a conventional microtiter well plate. Furthermore, the sandwich capture of sEVs with MBs in a liquid format and labelling with s-GNPs (E-NEXOS) or fluorescent reporters (O-NEXOS) has advantages over some of the emerging new technologies based on surface-based antibody capture of sEVs [17, 32, 33].

In liquid, mixing is promoted between the MBs and sEVs, whilst surface-based methods are limited by the lengthy diffusion of target analytes onto the surface of the chips where capturing antibodies are immobilised [34]. Furthermore, since the recognition of sEVs in these systems is performed by only one recognition site, it remains difficult to assess if the recognised target is in fact an sEV or a freely circulating protein.

E-NEXOS operates in a straightforward manner, with completion of the detection method occurring in minutes. This speed is achieved via the DEP step that results in the quick attraction of s-GNPs. Moreover, the additional activation step on E-NEXOS enables the unbiased detection of all s-GNPs trapped in the nanogaps (**Fig. 3F**). Further differentiation points of E-NEXOS compared to other similar techniques are: it requires no surface chemistry and no microfluidics; it has unlimited shelf-life; it is easy-to-use, and it is highly scalable. Additionally, the detection of sEVs with low abundant surface markers may be facilitated in E-NEXOS as it only requires a single binding to occur between an s-GNP and an sEV.

Regarding O-NEXOS, the method is high-throughput and easy to operate throughout, showing the highest sensitivity when compared to gold standards techniques in sEVs detection (**Fig. 7**).

Importantly, the two methods are ultra-sensitive and can be exploited to provide new dimensions in determining the concentration of TEVs, TEPs and epitopes per sEV (**Fig. 6**). To our knowledge, we show for the first time a methodology that is capable of this.

As our study predominantly concerns validation of the technology, the potential applications enabled by these dimensions are not explored. However, we propose several examples for future developments. For example, concerning the detection of disease biomarkers contained in sEVs, it may be important to quantify the number of disease epitopes to more accurately diagnose a certain indication or to stratify patients into different sub-classes of the disease [12, 35, 36]

Additionally, in drug discovery, determining the payload of molecules (e.g. proteins, antibodies) in engineered sEVs may validate or indicate that the established processes may require optimization [6]. Finally, these dimensions may enable the development of an atlas of sEV subpopulations based on their epitopes’ expression levels and make intelligent predictions on their content and origin [37].

The ability to provide such dimensions relies on the differences between E-NEXOS and O-NEXOS. Given the substantially smaller size of poly-HRP strep molecules (∼ 9.4 nm) vs s-GNPs (200 nm), and the similar size of poly-HRP strep and IgG antibodies (∼ 14.5 nm), we estimated that poly-HRP strep molecules can bind to most available target epitopes, if labeled with detection antibodies on the MBs-captured sEVs. Conversely, because the s-GNPs (∼ 200 nm) are twice the average size of sEVs (∼ 100 nm), they are limited in the number of MBs-captured sEVs they can bind to. Interestingly, we determined theoretically and experimentally that no more than 2 s-GNPs can bind per MB-captured sEV (**Fig. 5C** and **Table S2**).

We applied the methodology of detecting the TEVs from an enriched sample of CD9^+^CD81^+^ MCF-7 sEVs to the detection of CD9^+^CD81^+^ and CD9^+^HER2^+^ sEVs. We recognise that the sEVs are originated from different cell cultures and, most likely, display a different number of target epitope biomarkers per sEV. However, we noticed that the MBs used in this work have a very high capture efficiency (**Table S1** and **Fig. S3**) and that no more than 2 s-GNPs can bind per sEV, regardless of the number of epitopes per sEV. Thus, we assumed that a variation in sEV epitopes should not affect the method for the determination of TEVs.

The presented E-NEXOS method has not yet reached the sensitivity of O-NEXOS, however it is on par or surpasses the sensitivity of sEV gold standard detection techniques (**Fig. 7**).

E-NEXOS is currently presented as proof of concept with an extremely low density of electrode pairs in the sensing area: 36 sensors per nanochip, each with 3 nanogap electrode pairs. (**Fig. 3A** and **Fig. 3B**). Given the compatible fabrication of E-NEXOS with standard procedures in the semiconductor industry, the number of nanogap-based sensors can be increased to around 2,000,000 nanogap-based sensors per nanochip by integrating CMOS technology, as demonstrated by others [38-40].

Crucially, this will lead to significant improvements in sensitivity, precision, and dynamic range as well as throughput, by adapting the design to a nanochip-well format similar to a microtiter well plate. We expect that these improvements will result in near-absolute quantification of target sEVs and enable the achievement of single-sEV resolution.

To conclude, we presented NEXOS as a novel electro-optical technology for the ultrasensitive detection of sEVs which also determines new dimensions, enabling new research and clinical applications in sEV based diagnostics and therapeutics. At its present state, O-NEXOS can be readily applied for such applications with superior sensitivity compared to other gold standard sEV detection techniques. Currently, E-NEXOS already possesses high sensitivity but its indicated electronics miniaturisation from 36 to 2,000,000 nanogap-based sensors per nanochip allows for further significant and impactful improvements. Therefore, the next generation of E-NEXOS can achieve extremely high precision and sensitivity for the translation of sEV diagnostics into standard clinical practice.

## ACKNOWLEDGMENTS

The authors would like to thank Prof. Crispin Barnes for welcoming selected Mursla’s team members as Visiting Researchers in his research group at the Cavendish Laboratory of the University of Cambridge, to Prof. Andy Parker for supporting it and to Dr. Adrian Ionescu for his technical guidance. The authors would further like to thank Prof. Yutaka Majima and Dr. Victor Serdio for the initial advancements in the nanochip developments, to Prof. Dr. Bernd Giebel for revising the manuscript and Tobias Tertel for performing the IFC experiments and EverZom for performing the SP-IRIS experiments, to Dr Karin Müller from the Cambridge Advanced Imaging Centre for her support and assistance in the imaging of sEVs by TEM, to Thomas Mitchell and Jonathan Griffiths from Cavendish Laboratory for their support and services in EBL writing, to Mr Barry Shores from Cambridge University, Department of Physics for his support and services in electronics hardware design and interfacing, and finally to Prof. Andrea C. Ferrari of the Cambridge Graphene Centre and Sue Murkett from The Nanoscience Centre for allowing access to develop the nanochips. Figures 1, 4 and S1 were partially created with BioRender.com.

## MATERIALS AND METHODS

### Cell culture and sEVs preparation

Breast cancer cell lines MCF-7 (ATCC, catalogue nr HTB-22) and BT-474 (ATCC, catalogue nr HTB-20) were grown in growth medium Advanced-DMEM (Gibco), supplemented with 5 % (v/v) FBS (Gibco) and 4 mM glutaMAX (Gibco), in a humidifying atmosphere with 5 % CO2 and 37 °C.

After reaching 70 % confluency in T175 flaks (ThermoFisher), the cells were gently washed three times with PBS and were cultured in Advanced-DMEM FBS-free medium.

After 48h, the conditioned medium (CM) was collected and centrifuged for 5 min at 300 *g*, followed by another centrifugation of 10 min at 3000 *g* and 4°C to remove cellular debris and other large contaminants.

The CM was then filtered using 0.22 µm PES filters (FisherBrand) and concentrated 220 x down to 500 μL with Amicon Ultra-15 100K MWCO Centrifugal filters (Merck Millipore).

By size exclusion chromatography (SEC), the sEVs were finally isolated using Izon qEV original/70 nm columns (IZON), on an Automatic Fraction Collector (AFC), according to the supplier’s instructions.

The first eluted three fractions, corresponding to a total of 1.5 mL of elution volume, were collected, pooled together, aliquoted in volumes of 50 µL and stored at -80 °C until further use.

### Characterization of sEVs by Nanoparticle tracking analysis

Particle number and size distribution of MCF-7 and BT-474 and sEV preparations was determined by Nanoparticle Tracking Analysis (NTA) using a NanoSight NS300 system (Malvern) configured with a 488 nm laser and a high sensitivity scientific CMOS camera.

Samples were diluted in particle-free PBS (Gibco), to an acceptable concentration, according to the manufacturer’s recommendations. The samples were analysed under constant flow rate and 3 × 60 s videos were captured with a camera level of 12. Data was analysed using NTA 3.4 software with a detection threshold of 7.

### Western Blot analysis of sEV markers

MCF-7 and BT-474 sEV preparations were normalized to a concentration of 1E10 part/mL by dilution in particle-free PBS (ThermoFisher). Then, 100 µL of each sample was lysed in 1 x RIPA lysis buffer (ab156034, Abcam) supplemented with 1 x Halt™ Protease Inhibitor Cocktail (Thermo Fisher) and incubated for 1 hour on ice followed by 2 × 30 sec sonication in ice-cold water.

After, the samples were mixed with NuPAGE™ LDS Sample Buffer (4X) (Invitrogen) and NuPAGE™ Sample Reducing Agent (10X) (Invitrogen).

After boiling for 10 min at 70 °C, 37 µL per sample was loaded on a NuPAGE™ 4 to 12%, Bis-Tris, 1.0–1.5 mm, Mini Protein Gels (Invitrogen). Electrophoresis occurred at 150 V, using MES as the running buffer. Protein ladders (26634, Thermo Scientific orLC5925, Invitrogen) were run along with the sEV samples.

Transfer was performed on PVDF membranes (IB24002, Invitrogen) using an iBlot 2 Dry Blotting System (Invitrogen).

Using an iBind™ Western Device (Thermo Fisher) and following the supplier instructions, the membranes were incubated sequentially with iBind™ solution for blocking, primary antibodies and secondary antibody. The primary antibodies used were anti-HER2 (ab16901, Abcam) at 1:400, anti-TSG101 (ab83, Abcam) at 1:700, anti-CD63 (ab59479, Abcam) at 1:1000, anti-CD81 (orb506485, Biorbyt) at 1:1000 or anti-CD9 (ab58989, Abcam) at 1:1000. The secondary antibody used was a secondary antibody anti-mouse (HAF007, R&D Systems) at 1:250 dilution.

Finally, membranes were subsequently incubated with SuperSignal™ West Pico PLUS. Chemiluminescent Substrate (Thermo Scientific) and the bands were visualized and acquired on a GBOX Chemi XRQ (Syngene™).

### Bead-based Flow Cytometry analysis of sEV markers

MCF-7 and BT-474 sEVs were characterized by bead-based Flow Cytometry (FCM) to further validate the presence of CD63, CD81, CD9, and HER2 in the sEVs preparations. The sEVs were attached to 4 µm aldehyde/sulphate latex beads (Invitrogen) by the mixing of 8E8 sEVs with 4E6 beads in 200 μL of PBS for 20 min with continuous rotation.

The suspensions were diluted to 300 μL with PBS and left incubating overnight under agitation. The reactions were stopped with 100 mM glycine in PBS, during 1 h and under continuous agitation. Bead-bound sEVs were centrifuged for 5 min at 12000 *g* and the PBS was replaced once. The bead-bound sEVs were centrifuged again for 5 min at 12000 *g*, supernatant was discarded and bead-bound sEVs were blocked by resuspension of the beads in 300 µL of PBS with 5% BSA. After 45 min of agitation, the beads were washed a second time in 2 % BSA and centrifuged once more for 5 min at 12000 *g*, and resuspended with primary antibodies either targeting CD63, CD81, CD9 or HER2, in 2 % BSA for 1 h with rotation. Samples were washed three times in 200 μl of 2% BSA and incubation with secondary antibody was done in the dark for 30 min, washed as previously described and resuspended in 600 µL 2 % BSA.

For tetraspanins detection, the mouse antibodies anti-CD63 (ab59479, Abcam), anti-CD81 (ab59477, Abcam) or anti-CD9 (ab58989, Abcam) were used. HER2 was detected with a recombinant human biotinylated antibody (FAB9589B, RnDSystems). Mouse isotype antibody (ab170190, Abcam) or human isotype biotinylated (NBP1-96855, Novus Biological) were used as controls. Secondary antibody anti-mouse AF488 (ab150113, Abcam) or PE-streptavidin (ab239759, Abcam) were used as reporters.

Analysis was performed by FCM using an Invitrogen Attune NxT. The channel YL1 (excitation 561 nm and emission 585/16 nm) or BL1 (excitation 488 nm and emission 530/30nm) were selected for detection of the PE or AF488 fluorophores, respectively. Results were visualized using Invitrogen Attune NxT Software. Single sEVs-beads complexes were gated using forward-scattered light (FSC) Area (A) (x-axis) and side-scattered light (SSC) Area (A) (y-axis). Data analysis was performed with FlowJo Software v10.6.1. The percentage of positive beads was calculated relative to the total number of beads analysed per sample (10,000 events). This percentage was therein referred to as the percentage of beads with sEV markers.

### sEVs characterization by Transmission Electron Microscopy

10 μL of freshly prepared sEVs at a concentration of 1E9 part/mL were spotted on a glow-discharged carbon-coated 400 mesh copper TEM grid and incubated for 2 minutes. To remove excess PBS, the TEM grid was washed in 2 consecutive 10 μL drops of deionized water for 30s each. Then, the grid was stained with 10 μL of uranyl acetate for 60 s. The grid was dried by tapping it on dust-free clean room paper. Finally, the grid was transferred to a FEI Tecnai G2 TEM and imaged at room temperature and 200 kV. Images were acquired with a bottom-mounted AMT CCD camera, using a bright field imaging mode.

### Cell-culture derived sEVs preparation for detection using NEXOS and gold standard technologies

MCF-7 and BT-474-derived sEVs were prepared as previously described. Then, 50 µL sEV aliquots were gently thawed and the concentration of sEVs in the aliquots was determined by NTA as before. Both the MCF-7 and BT-474 sEVs used in this study, originated from 2 batches of sEV preparations each.

The sEVs were serially diluted to concentrations in the range of 5E8 and 7.5E5 part/mL, up to four samples for each concentration point and totaling 200 µL for measurement with E-NEXOS and O-NEXOS. The sEVs used for detection with the additional technologies were prepared identically and originated from the same batches of sEV preparations.

### Processing of human plasma and spiking with sEVs for detection in NEXOS

4 mL fresh human plasma derived from whole blood in K2EDTA Vacutainers was acquired (Cambridge bioscience). The human plasma was ethically consented and originated from a pool of healthy, paid volunteers (Research Donors).

The human plasma was slowly thawed on ice and divided into two portions of 2 mL each. One of the portions was spiked with BT-474 sEVs, reaching to a final concentration of 1.67E9 part/mL. Then, both the spiked and non-spiked samples were treated equally by a centrifugation step of 15 min, at 3000 *g* and at 4°C followed by another centrifugation of 20 min at 10000 *g* and 4 °C to remove any cell debris, large particles or aggregates. Then, the samples were filtered using a 0.22 µm PES filter and 1.8 mL of each sample was processed by SEC using an Izon qEV2 70 nm column on an AFC. As before, the first eluted three fraction of each sample, and corresponding to a total of 5.4 mL per sample, were collected and pooled together. Considering the dilution step in the process with SEC, the concentration of spiked BT-474 sEVs was now of 5.57E8 part/mL in maximum.

Finally, we serially diluted the processed spiked plasma to samples containing BT-474 sEVs concentrations of 1E8, 5E7, 2.5E7, 1E7, 5E6 and 2.5E6 by addition of non-spiked processed plasma, totalling a final volume of 200 µL per sample, each concentration point with n = 4. These samples were used to determine the sensitivity of NEXOS in detecting sEVs spiked in processed plasma as described below.

### E-NEXOS nanochip design and fabrication

The nanochips were designed on AutoCAD and fabricated on n-doped silicon 4’’ wafers of 525 µm thickness, coated with 100 nm WET thermal oxide. The design included 37 dyes, each one corresponding to one E-NEXOS nanochip, with a separation of 360 µm between each dye. The fabrication was divided into two layers, each with a lithography and metal deposition steps. The nanogaps were fabricated in the first layer. First, 100 nm thick PMMA 950K A4 positive resist was spin-coated on the substrate. Then, the resist was exposed by electron beam lithography at a dose of 590 µC/cm^2^ (Vistec VB6) and developed in a solution made of IPA:MIBK:MEK 15:5:1. After development, Ti 5/Au 30 (thicknesses in nm) were deposited by electron-beam evaporation (Lesker e-beam evaporator PVD 200 Pro) and the gold-patterned nanogaps were revealed by lift-off in acetone. The contact lines and pads were fabricated in the second layer. AZ5214-E 1000 nm thick image reversal photoresist was spin-coated on the substrate. The pattern was exposed and defined by photolithography (SUSS Microtech MA/BA6 mask aligner) with critical dimension < 2 μm. The photoresist layer was developed in an aqueous solution of H20:AZ351B 4:1. After development, Ti 5/Au 30 (in nm) layers were deposited, again, by evaporation using the Lesker PVD 200 Pro, followed by lift-off using AZ100.

Finally, the wafers were diced into 37 nanochips using a mechanical dicer (DISCO DAD341) and the nanogaps were quality-checked by room temperature SEM (FEI Magellan 400L) ETD detector at 5 KV, with secondary electrons imaging, to confirm the physical separation and dimensions of the structures. Additionally, after using E-NEXOS, the presence of s-GNPs on selected sensors was also visualized by SEM.

### Labelling of sEVs and mixing with MBs

sEVs were thawed on ice and diluted to target sEV concentrations, as previously explained, in a final buffer composition of 1 x PBS, 0.1 % BSA, 0.02 % Tween®20. 100 ng of detection antibodies anti-CD81 (ab239238, Abcam) or anti-HER2 (FAB9589B-100, R&D Systems) were added to each sEV sample totalling a final total volume of 200 µL per well in a U-shaped 96-well plate and incubated for 45 minutes at RT and 200 RPM. Magnetic beads (MBs) (Dynabeads® M-270 Epoxy beads), were coupled to sEVs capture antibodies anti-CD9 (ab58989, Abcam), anti-CD81 (orb506485, Biorbyt) or isotype IgG (orb343761, Biorbyt), by using the Dynabeads™ Antibody Coupling Kit whilst following the supplier’s instructions. Briefly, the MBs were washed as per the kit instructions and incubated ON at 37 ºC, under constant rotation in LoBind microcentrifuge tubes (Eppendorf). The amount of capture antibody used in the reactions was 10 µg per mg of MBs. Tween®20 (0.05 %) was added in HB and LB wash buffers for improved stringency as recommended by the supplier. Washings of the MBs was performed with a DynaMag – Spin Magnet (Invitrogen).

7.5 µL or 15 µL of MBs, were added to the sEVs diluted in PBS or sEVs spiked in processed plasma, respectively. The capture of sEVs by the MBs occurred for 3 h with a step of up-and-down pipetting every 30 min to guarantee the homogenous mixing of the MBs and the sEVs with the aid of an automatic pipetting robotic (Opentrons). After, the MBs were washed three times in PBS with 0.05 % Tween®20 and 0.1 % BSA before proceeding for detection in E-NEXOS or O-NEXOS.

### Determination of CD9^+^ sEVs capture rate and enrichment of CD9^+^CD81^+^ sEVs

To determine the capture rate of sEVs by the MBs, MCF-7 sEVs were captured with MBs pre-coated with antibody anti-CD9 as explained before. As opposed to the prior methods, a detection antibody was not used to pre-label the sEVs. After an incubation period of 3 h, the MBs-bound sEVs were washed in a magnetic field, buffer was removed and the sEVs were eluted from the MBs in 0.2 M Glycine, pH 2.6 0.05 % Tween®20 (equal v/v Glycine solution to MBs). The pH was restored to pH 7.2 with 1 M Tris-HCL, pH 9.0 (1 µL of TRIS per 10 µL of Glycine). Then, the concentration of sEVs was determined by NTA. Enriched CD9^+^ sEVs, at a concentration of 7.29E9 part/mL, were captured again with MBs anti-CD9. The concentration of sEVs remaining in the supernatant was determined by NTA.

To specifically enrich for CD9^+^CD81^+^ sEVs, MCF-7 sEVs, were first captured with MBs anti-CD9 and eluted in glycine solution as previously, and captured again with MBs anti-CD81 and eluted once again in glycine solution. Then, we serially diluted the enriched CD9^+^CD81^+^ sEVs to samples starting at a concentration of 1E8 part/mL down to 1E6 part/mL and used E-NEXOS as explained after in low sensitive and high sensitive modes to determine the number of bound s-GNPs per target sEV.

### Dielectrophoresis modelling in E-NEXOS

We designed a 3D finite element model using COMSOL Multiphysics to determine the electric field characteristics around the nanogap and the dieletrophoresis (DEP) force exerted on a double-shell gold nanoparticle simulating the s-GNPs used in our experiments.

The model comprised a 12 µm × 12 µm × 8 µm simulation block with 2 gold electrodes of 300 nm width and 35 nm thickness separated by 50 nm and sitting on top of a SiO2 substrate. The model considered the effective complex Clausius-Mossoti factor of a s-GNP with a gold core (200 nm) and 2 shell layers (thiol and streptavidin), suspended in deionized water.

The cross-sectional gradient of the electric field intensity was simulated over the 3 orthogonal planes across the nanogap using COMSOL’s Electrostatics module. We fixed 0 V in one of the electrodes, and the quadratic mean of 2 V in the other electrode (as we used experimentally).

### E-NEXOS calibration, sEVs detection and determination of TEVs

The concentration of customized 200 nm streptavidin GNPs (s-GNPs) (Cytodiagnostics) was determined by NTA. Then, a suspension with s-GNPs at a concentration of 2E9 part/mL was serially washed thrice by centrifugation for 15 min at 400 *g* and resuspended in either 1 % PBS (high range sensitivity) or ultra-pure water (low range sensitivity). The s-GNPs where the buffer was exchanged to 1 % PBS, were serially diluted to concentrations ranging from 1E8 part/mL down to 1E6 part/mL by the addition of 1 % PBS in a volume of 200 µL per sample. Otherwise, the s-GNPs where the buffer was exchanged to Ultra-pure water, were serially diluted to concentrations ranging from 2.5E6 part/mL down to 7E4 part/mL by addition of ultra-pure water, again, in a volume of 200 µL per sample. After, 45 µL of each sample were pipetted onto the E-NEXOS nanochips and corresponding signal was acquired as explained below. A calibration curve was derived, allowing to correlate the number of hits with the concentration of s-GNPs in solution. Samples with serially diluted sEV concentrations were captured with MBs anti-CD9 and sequentially washed in 1 x PBS, 0.1 % BSA, 0.02 % Tween®20 under a magnetic field. The buffer was removed and 200 µL of s-GNPs at a concentration of 2E9 part/mL were added to the complexes formed of MBs-bound sEVs and incubated for 30 min under constant agitation. The complexes were washed and the GNPs were eluted in Glycine, pH 2.6 0.05 % Tween®20 (equal v/v Glycine solution to MBs). The pH was restored to pH 7.2 and then exchanged to either 1 % PBS or ultra-pure water. Finally, 45 µL of each sample were pipetted onto the E-NEXOS nanochips and the corresponding signals were acquired as explained below. For the determination of TEVs, enriched CD9^+^CD81^+^ sEVs, prepared as explained above, were labelled again with detection antibody anti-CD81 for 45 min and captured with MBs anti-CD9 for 3 h. MBs-bound sEVs were washed and s-GNPs added and incubated as above. The GNPs were eluted and buffer was exchanged to either 1 % PBS or ultra-pure water. Finally, 45 µL of each sample were pipetted onto the E-NEXOS nanochips and the corresponding signals were acquired. A calibration curve was derived, allowing to correlate the concentration of s-GNPs with the concentration of target sEVs in solution.

### E-NEXOS nanochip interface and signal acquisition

The nanochips were interfaced with a dedicated readout system designed and developed by TronicsZone. The interface occurred via a plug-in printed circuit board (PCB) developed by Newbury Electronics Ltd, which connects the E-NEXOS nanochip and the readout system via spring-loaded pins. The readout was used for sourcing DC/AC to the nanochips and for measuring DC currents after the trapping of s-GNPs on the nanogaps.

The sensing area of the nanochips was protected from exterior wiring or contact with the PCB with barriers made of elastosil E-41 (Wacker).

For signal acquisition, several steps were performed. First, sequential IV curves on every sensor (0-5V), defined the sensors baseline. Sensors with current leakages above 50 nA were excluded. Then, 45 μL of s-GNPs suspensions, prepared as explained before, were pipetted over the sensing area and DEP was performed at 2 V AC and 100 kHz for 60s in every sensor.

After DEP, the solution was blown-dried with pressurized filtered air. Subsequently, we applied an activation step of 5 V for 20s, followed by an IV curve (0-5V), sequentially in every sensor. The sensors with current readings under 1 µA were counted as disconnected. Data analysis was automated with a script developed in Python.

### O-NEXOS calibration, sEVs detection and determination of TEPs

Pierce™ Streptavidin Poly-HRP (Poly-HRP strep) (Thermo Scientific) was diluted 1:6000 in 1 x PBS, 0.1% BSA, 0.02 % Tween®20, corresponding to 9.12E11 streptavidin molecules/mL, and then serially diluted into several lower concentrations. 10 µL of each concentration was transferred into a U-shaped 96-well plate and incubated with 100 µL QuantaBlu™ Fluorogenic Peroxidase Substrate Kit (Thermo Scientific™) for 40 minutes along with the sEV samples to be detected on O-NEXOS. The reactions were stopped by adding 100 µL of stop solution and 190 µL of each sample was transferred to a F-shaped 96-well plate. Fluorescence signals were obtained at Ex/Em = 320/405 nm with a Fluoroskan Ascent FL Microplate Fluorometer (Thermo Scientific).

A calibration curve was derived, allowing to correlate the fluorescence signal with the concentration of HRP-strep molecules in solution.

Samples with serially diluted sEV concentrations were captured with MBs anti-CD9 and sequentially washed. The buffer was removed and 200 µL of Poly-HRP strep 1:6000 diluted was incubated with the complexes for 30 min. After washing, as in O-NEXOS calibration, 100 µL QuantaBlu™ were added and incubated for 40 minutes. The reactions were stopped with 100 µL of stop solution and 190 µL of each sample was transferred to a F-shaped 96-well plate for signal measurement at Ex/Em = 320/405 nm.

### ELISA

The colorimetric ExoELISA-ULTRA Complete Kit (CD81 Detection) (Systems Biosciences) was used for CD81 detection of BT-474 sEVs. A standard curve and the ELISA procedure were performed according to the supplier’s instructions.

## REFERENCES

1. Mathieu, M., et al., Specificities of secretion and uptake of exosomes and other extracellular vesicles for cell-to-cell communication. Nature Cell Biology, 2019. 21(1): p. 9–17.

2. Kalluri, R. and V.S. LeBleu, The biology, function, and biomedical applications of exosomes. Science, 2020. 367(6478).

3. Yates, A.G., et al., In sickness and in health: The functional role of extracellular vesicles in physiology and pathology in vivo - PART I. Journal of Extracellular Vesicles, 2022. 11(1): p. e12151.

4. Yates, A.G., et al., In sickness and in health: The functional role of extracellular vesicles in physiology and pathology in vivo - PART II. Journal of Extracellular Vesicles, 2022. 11(1): p. e12190.

5. Pink, R.C., et al., Utilising extracellular vesicles for early cancer diagnostics: benefits, challenges and recommendations for the future. Br J Cancer, 2022. 126(3): p. 323–330.

6. Herrmann, I.K., M.J.A. Wood, and G. Fuhrmann, Extracellular vesicles as a next-generation drug delivery platform. Nature Nanotechnology, 2021. 16(7): p. 748–759.

7. van Niel, G., G. D’Angelo, and G. Raposo, Shedding light on the cell biology of extracellular vesicles. Nature Reviews Molecular Cell Biology, 2018. 19(4): p. 213–228.

8. Garcia-Martin, R., et al., Tissue differences in the exosomal/small extracellular vesicle proteome and their potential as indicators of altered tissue metabolism. Cell Rep, 2022. 38(3): p. 110277.

9. Becker, A., et al., Extracellular Vesicles in Cancer: Cell-to-Cell Mediators of Metastasis. Cancer cell, 2016. 30(6): p. 836–848.

10. Kosaka, N., et al., Exploiting the message from cancer: the diagnostic value of extracellular vesicles for clinical applications. Experimental & Molecular Medicine, 2019. 51(3): p. 1–9.

11. Margolis, E., et al., Predicting high-grade prostate cancer at initial biopsy: clinical performance of the ExoDx (EPI) Prostate Intelliscore test in three independent prospective studies. Prostate Cancer and Prostatic Diseases, 2021.

12. Hoshino, A., et al., Extracellular Vesicle and Particle Biomarkers Define Multiple Human Cancers. Cell, 2020. 182(4): p. 1044-1061.e18.

13. Ferguson, S. and R. Weissleder, Modeling EV Kinetics for Use in Early Cancer Detection. Advanced Biosystems, 2020. 4(12): p. 1900305.

14. Arab, T., et al., Characterization of extracellular vesicles and synthetic nanoparticles with four orthogonal single-particle analysis platforms. Journal of Extracellular Vesicles, 2021. 10(6): p. e12079.

15. Jahani, Y., et al., Imaging-based spectrometer-less optofluidic biosensors based on dielectric metasurfaces for detecting extracellular vesicles. Nature Communications, 2021. 12(1): p. 3246.

16. Park, J., et al., An integrated magneto-electrochemical device for the rapid profiling of tumour extracellular vesicles from blood plasma. Nature Biomedical Engineering, 2021. 5(7): p. 678–689.

17. Kilic, T., et al., Multielectrode Spectroscopy Enables Rapid and Sensitive Molecular Profiling of Extracellular Vesicles. ACS Central Science, 2022. 8(1): p. 110–117.

18. Liu, C., et al., Low-cost thermophoretic profiling of extracellular-vesicle surface proteins for the early detection and classification of cancers. Nature Biomedical Engineering, 2019. 3(3): p. 183–193.

19. Vaz, R., et al., Breaking the classics: Next-generation biosensors for the isolation, profiling and detection of extracellular vesicles. Biosensors and Bioelectronics: X, 2022. 10: p. 100115.

20. Xu, L., et al., Development of a simple, sensitive and selective colorimetric aptasensor for the detection of cancer-derived exosomes. Biosensors and Bioelectronics, 2020. 169: p. 112576.

21. Nasibeh Karimi, R.D., Tomás Dias, Jan Lötvall, Cecilia Lässer, Tetraspanins distinguish separate extracellular vesicle subpopulations in human serum and plasma - Contributions of platelet extracellular vesicles in plasma samples. J. Extracellular vesicles, 2022.

22. Dash, M., et al., Exosomes isolated from two different cell lines using three different isolation techniques show variation in physical and molecular characteristics. Biochimica et Biophysica Acta (BBA) - Biomembranes, 2021. 1863(2): p. 183490.

23. Xiong, H., et al., Recent Progress in Detection and Profiling of Cancer Cell-Derived Exosomes. Small, 2021. 17(35): p. 2007971.

24. Green, T.M., et al., Breast Cancer-Derived Extracellular Vesicles: Characterization and Contribution to the Metastatic Phenotype. BioMed research international, 2015. 2015: p. 634865–634865.

25. Shin, I., HER2 Signaling in Breast Cancer, in Translational Research in Breast Cancer, D.-Y. Noh, W. Han, and M. Toi, Editors. 2021, Springer Singapore: Singapore. p. 53–79.

26. Ménard, S., et al., Biologic and therapeutic role of HER2 in cancer. Oncogene, 2003. 22(42): p. 6570–6578.

27. Théry, C., et al., Minimal information for studies of extracellular vesicles 2018 (MISEV2018): a position statement of the International Society for Extracellular Vesicles and update of the MISEV2014 guidelines. J Extracell Vesicles, 2018. 7(1): p. 1535750.

28. Victor Serdio, T.D. Pierre Arsène, GB2583550 - Biosensor conditioning method and system. 2020: UK.

29. Tomás Dias, P.A., GB2595423 - Biosensor conditioning method and system. 2021: UK.

30. Mustapic, M., et al., Plasma Extracellular Vesicles Enriched for Neuronal Origin: A Potential Window into Brain Pathologic Processes. Frontiers in Neuroscience, 2017. 11.

31. Popovic, M., et al., Isolation of anti-extra-cellular vesicle single-domain antibodies by direct panning on vesicle-enriched fractions. Microbial Cell Factories, 2018. 17(1): p. 6.

32. Hajian, R., et al., Rapid and Electronic Identification and Quantification of Age-Specific Circulating Exosomes via Biologically Activated Graphene Transistors. Advanced Biology, 2021. 5(7): p. 2000594.

33. Qiu, G., et al., Detection of Glioma-Derived Exosomes with the Biotinylated Antibody-Functionalized Titanium Nitride Plasmonic Biosensor. Advanced Functional Materials, 2019. 29(9): p. 1806761.

34. Squires, T.M., R.J. Messinger, and S.R. Manalis, Making it stick: convection, reaction and diffusion in surface-based biosensors. Nat Biotechnol, 2008. 26(4): p. 417–26.

35. Rontogianni, S., et al., Proteomic profiling of extracellular vesicles allows for human breast cancer subtyping. Communications Biology, 2019. 2(1): p. 325.

36. Trino, S., et al., Clinical relevance of extracellular vesicles in hematological neoplasms: from liquid biopsy to cell biopsy. Leukemia, 2021. 35(3): p. 661–678.

37. Xiao, X., et al., Intelligent Probabilistic System for Digital Tracing Cellular Origin of Individual Clinical Extracellular Vesicles. Analytical Chemistry, 2021. 93(29): p. 10343–10350.

38. Fuller Carl, W., et al., Molecular electronics sensors on a scalable semiconductor chip: A platform for single-molecule measurement of binding kinetics and enzyme activity. Proceedings of the National Academy of Sciences, 2022. 119(5): p. e2112812119.

39. Hall, D.A., et al. 16.1 A nanogap transducer array on 32nm CMOS for electrochemical DNA sequencing. in 2016 IEEE International Solid-State Circuits Conference (ISSCC). 2016.

40. Huang, M., et al. Large-scale plasmonic microarray: A new approach for label-free high-throughput biosensing and screening. in 2012 Conference on Lasers and Electro-Optics (CLEO). 2012.

