## Supplementary Information for "An electro-optical bead-nanochip technology for the ultrasensitive and multi-dimensional detection of small extracellular vesicles and their markers"

### TABLE OF CONTENTS

1 – Supplementary theory and sensing principle of E-NEXOS

2 – E-NEXOS method optimization

3 – Optimization and validation of sEVs capture with MBs

4 – Ratio of s-GNPs to sEVs and determination of TEVs and TEPs

5 – Detection of pre-prepared cancer cell-derived sEVs with NEXOS technology

6 – Detection of sEVs with different technologies

### 1 – Supplementary theory and sensing principle of E-NEXOS

In brief, E-NEXOS is a patented nanoelectronics approach for the sensitive detection of nanoparticles in fluids. E-NEXOS is enabled by nanochips comprising nanogap-based sensors, a readout for signal generation and acquisition and a PCB for interfacing the nanochip with the readout.

The nanochip is composed of multiple sensors, each one being able to attract metallic nanoparticles via a "sink effect" and sense the nanoparticles trapped in the nanogap via consecutive steps of activation and sensing. The sensors are arranged as electrode pairs separated by nanogaps of 50 nm.

The attraction of nanoparticles results from the application of an AC voltage across the electrodes pair, which generates a non-uniform electric field perpendicular to the nanogaps. This field polarizes conductive nanoparticles in the vicinity with a positive dielectrophoresis (DEP) force and is described by equation S1:

$$F_{DEP} = 2\pi\varepsilon_m R^3 Re\{CM(\omega)\}\nabla\left|\vec{E}_{rms}\right|^2 \quad \text{(S1)}$$

where for a particle of radius R in a solution, with a dielectric permittivity $\varepsilon_m$ , the $\vec{E}_{rms}$ is the root mean square value of the electric field gradient. The frequency-dependent complex Clausius–Mossotti factor, $CM(\omega)$, is given by equation S2:

$$CM(\omega) = \frac{\varepsilon_p{}^* - \varepsilon_m{}^*}{\varepsilon_p{}^* + 2\varepsilon_m{}^*} \quad \text{(S2)}$$

where the $\varepsilon_p{}^*$ and $\varepsilon_m{}^*$ are the complex permittivity of the particle and the medium, respectively.

Attracted nanoparticles land and remain trapped on the nanogaps, connecting therefore the electrodes pair.

Once connected, an activation step with 5 V DC, fuses the electrodes and the nanoparticle together, allowing to discriminate, in the final sensing step, the sensors that are connected by at least a GNP from the sensors that are disconnected. The signal amplitude shifts from picoamperes (open circuit) to milliamperes (closed circuit) at least, a nine order of magnitude increase and unbiased detection of even a single GNP is obtained (**Fig. S1**).

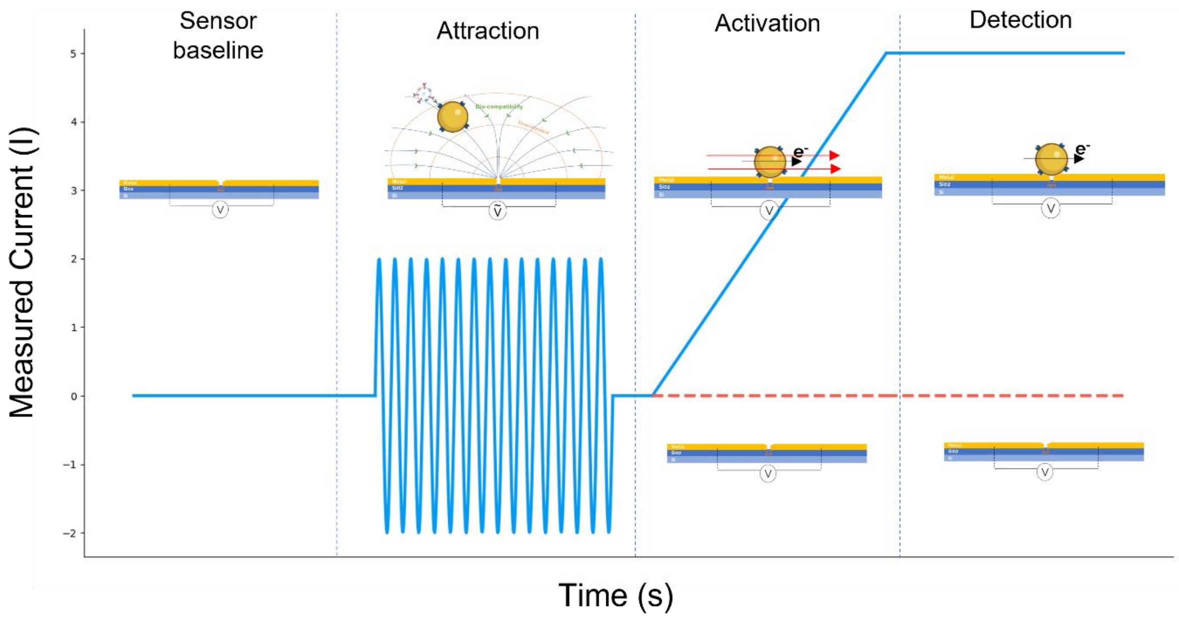

**Fig. S1. Schematics of the attraction and sensing steps in two scenarios: nanogap is bridged or not bridged by a s-GNP.** Sensor baseline: Initially, a sensor baseline is established, confirming whether the nanogap is unbridged (open circuit). Attraction: Next, AC voltage is applied which polarizes and attracts the target s-GNPs towards the nanogap. Activation: DC voltage is applied across the s-GNP-bridged nanogap electrodes, resulting in the fusion of the s-GNP with the electrodes pairs (closed circuit). Detection: High current generated from the s-GNP fused with the nanogap electrodes can be unbiasedly detected. In unbridged nanogaps, no increase in current is detected at the nanogap-based sensors.

**2 – E-NEXOS method optimization**

For a given nanochip, our method offers tuneable sensitivity and detection ranges by the selection of different parameters that together result in different $F_{DEP}$ over a nanoparticle, in the vicinity. These can include the size of streptavidin-GNPs (s-GNPs), AC field for sensor bias, permittivity of the media in which the s-GNPs are eluted or the attraction surface area amongst others.

Because $F_{DEP}$ varies proportionally to the cubic of a s-GNP radius ($R^3$), it is expected that the larger a s-GNP is, the easier it is to attract it to the nanogap. However, the drag force (resistance to motion) and force of gravity are also predominant in larger nanoparticles. Therefore, we compared biocompatible s-GNPs of 100 and 200 nm, diluted in ultra-pure water, at different concentrations and measured the percentage of hits (%) for a fixed AC bias of 2 V.

We observed that for the different concentration of s-GNPs, the percentage of hits (%) increases proportionally with the concentration of s-GNPs but that a larger number of hits is consistently obtained with the 200 nm s-GNPs. Furthermore, we reached a LOD of 7.5E4 s-GNPs/mL offering 6.7x more sensitivity than with s-GNPs of 100 nm (**Fig. S2-A**).

$F_{DEP}$ is also proportional to the gradient of the squared electric field intensity and the electric field intensity is described, in its simplest form, by equation S3:

$$E = V/d \tag{S3}$$

Where V is the electric field potential and d is the nanogap separation between the electrodes.

We compared the performance of the method at AC bias of 2 V, as before, with that of 4 V and 0.8 V. As predicted, 0.8 V offered the lowest sensitivity and operation at higher concentrations ranges (**Fig. S2-B**). The highest sensitivity was obtained at 4 V AC with a LOD of 2.5E4 part/mL but we determined that electric fields generated by more than 2 V AC were more likely to compromise the structure of the nanogaps and resulted in false positive nanogap connections (data not shown).

Additionally, we compared media with different permittivity. Again, we serially diluted 200 nm s-GNPs, either in ultra-pure water or in 1 % PBS and, for different concentrations, we measured the number of hits (%) for a fixed AC bias of 2 V. We observed that the highest sensitivity is obtained when the s-GNPs were eluted in ultra-pure water (LOD of 7.5E4 part/mL) when compared to 1 % PBS (LOD of 2.5E6 part/mL) (**Fig. S2-C**). Additionally, we concluded that simply by eluting the s-GNPs in the two different media, we could scan a large range of s-GNPs concentrations, from 7.5E4 to 1E8 part/mL (**Fig. 3-G**) and therefore detect a large range of sEV concentrations.

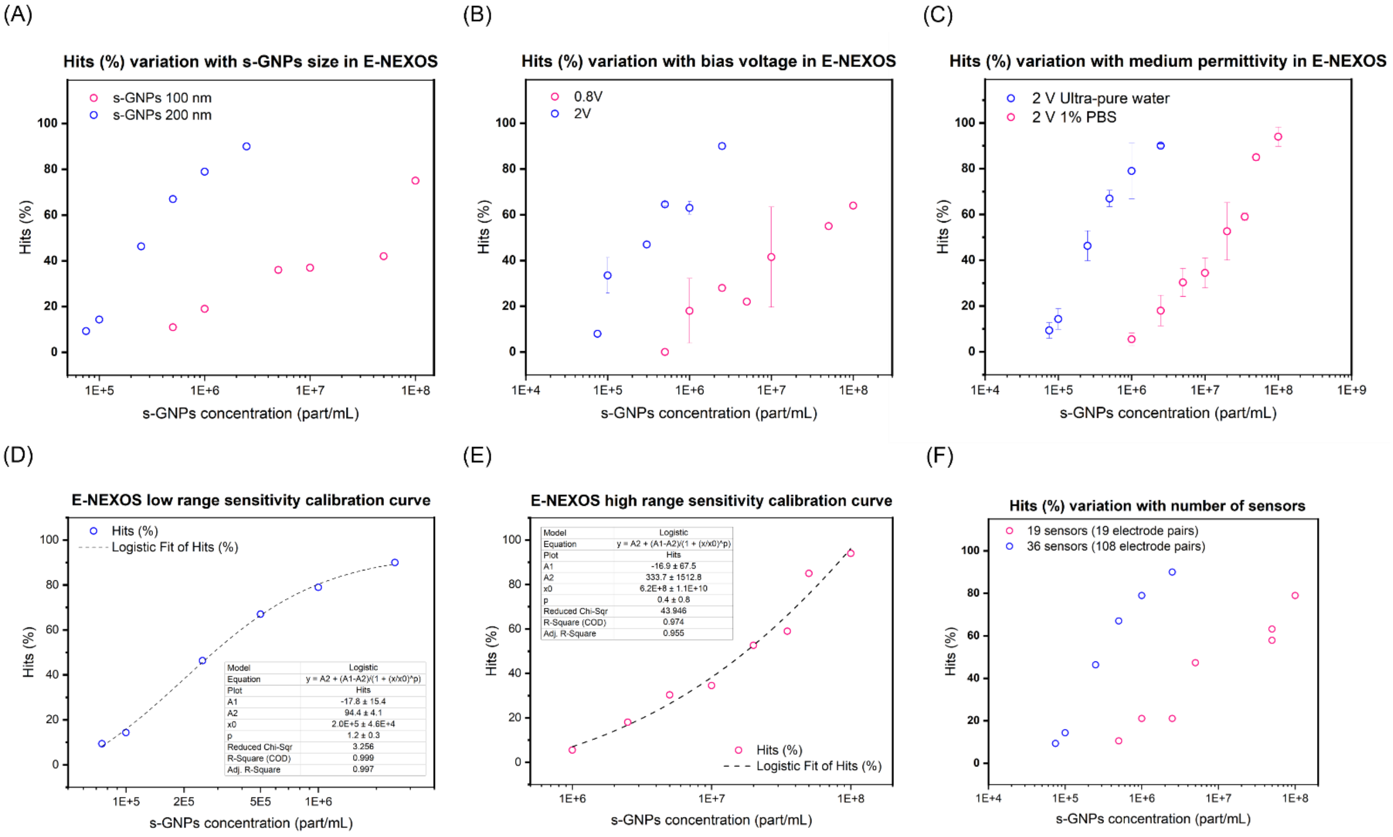

**Fig. S2. E-NEXOS optimization parameters and high range and low range calibration curves.** (A) Hits (%) in E-NEXOS in function of the size of s-GNPs at serially diluted s-GNP concentrations. (B) Hits (%) in E-NEXOS in function of the biasing voltage for nanogap attraction of s-GNPs at serially diluted s-GNP concentrations. (C) Hits (%) in E-NEXOS in function of s-GNPs concentrations and by diluting the s-GNPs in media with different permittivity, namely ultra-pure water and 1 % PBS. (D) E-NEXOS high range calibration curve as a percentage of hits (%) in function with the concentration of s-GNPs. (E) E-NEXOS low range calibration curve as a percentage of hits (%) in function with the concentration of s-GNPs. (F) Dynamic range comparison of E-NEXOS nanochips with 19 (19 electrode pairs) and 36 (108 electrode pairs) sensors.

The calibration equations fit in a logistic model as demonstrated (**Fig. S2-D** and **S2-E**).

Lastly, we compared the sensitivity of chips with 36 sensors (108 electrode pairs) against chips with 19 sensors (19 electrode pairs) for a fixed size of GNPs and identical electrical parameters. We observed that with the increased nanogaps sensor area (36 sensors, with 3 electrode pairs each), the sensitivity achieved is of 7.5E4 whereas for chips with 19 sensors, with 1 electrode pair each, the sensitivity is limited to 4.5E5 part/mL (**Fig. S2-F**).

**3 – Optimization and validation of sEVs capture with MBs**

We developed a bead-based capture method where 2.8 µm Magnetic Beads (MBs) were pre-coated with capture antibodies and used after to capture and isolate sEVs from biofluids. Briefly, we coupled 10 µg antibody per 1 mg Dynabeads (antibody in excess of the recommended amount) and determined the amount of antibodies efficiently coupled to the MBs with a BCA protein assay.
The results are shown as the mean of different preparations for the antibodies anti-CD9, anti-CD81, IgG isotype, utilized in our assays, with the background of MBs (without antibody) subtracted (**Fig. S3-A**). We observed a good correlation with the coating values estimated by the supplier.
Additionally, we determined the capture rate of target MCF-7 sEVs at different timepoints, with MBs pre-coated with anti-CD9 or with isotype IgG (used as control).
MBs pre-coated with anti-CD9 antibody or with isotype IgG antibody (used as control), were mixed with MCF-7 sEVs prepared by differential centrifugation, filtration and SEC as explained in Materials and Methods, and we assessed different periods of incubation between the MBs and the sEVs, namely 1 h, 3 h and 16 h for an initial concentration of the sEVs of 1.3E9 part/mL (or a quantity of 2.8E8 sEVs) per reaction as determined by NTA.
After incubation, we determined by NTA the concentration of sEVs remaining in the supernatant at the different timepoints. Knowing the initial concentration in each sample, we then calculated the concentration of captured sEVs for each timepoint and determined the capture rate of the method.
We observed that the best signal-to-noise ratio is achieved at 3 h incubation and, thus, we selected 3 h for the capture of sEVs in our assays (**Fig. S3-B**).
Next, we validated the efficiency of our method in capturing an enriched population of $CD9^+$ sEVs. For that, we prepared MCF-7 sEVs by differential centrifugation, filtration and SEC as previously explained. Then, we captured $CD9^+$ sEVs with MBs anti-CD9. The MBs-bound sEVs were washed in a magnetic field and the elution of the $CD9^+$ sEVs was performed in glycine at pH 2.6, as per previously described and validated methods. Finally, pH was restored to 7.2 (See Materials and Methods). When serially capturing and eluting sEVs with isotype IgG beads, no sEVs were enriched as determined by NTA suggesting that any unspecific adsorption of sEVs to beads is detached during our washing protocol (data not shown).

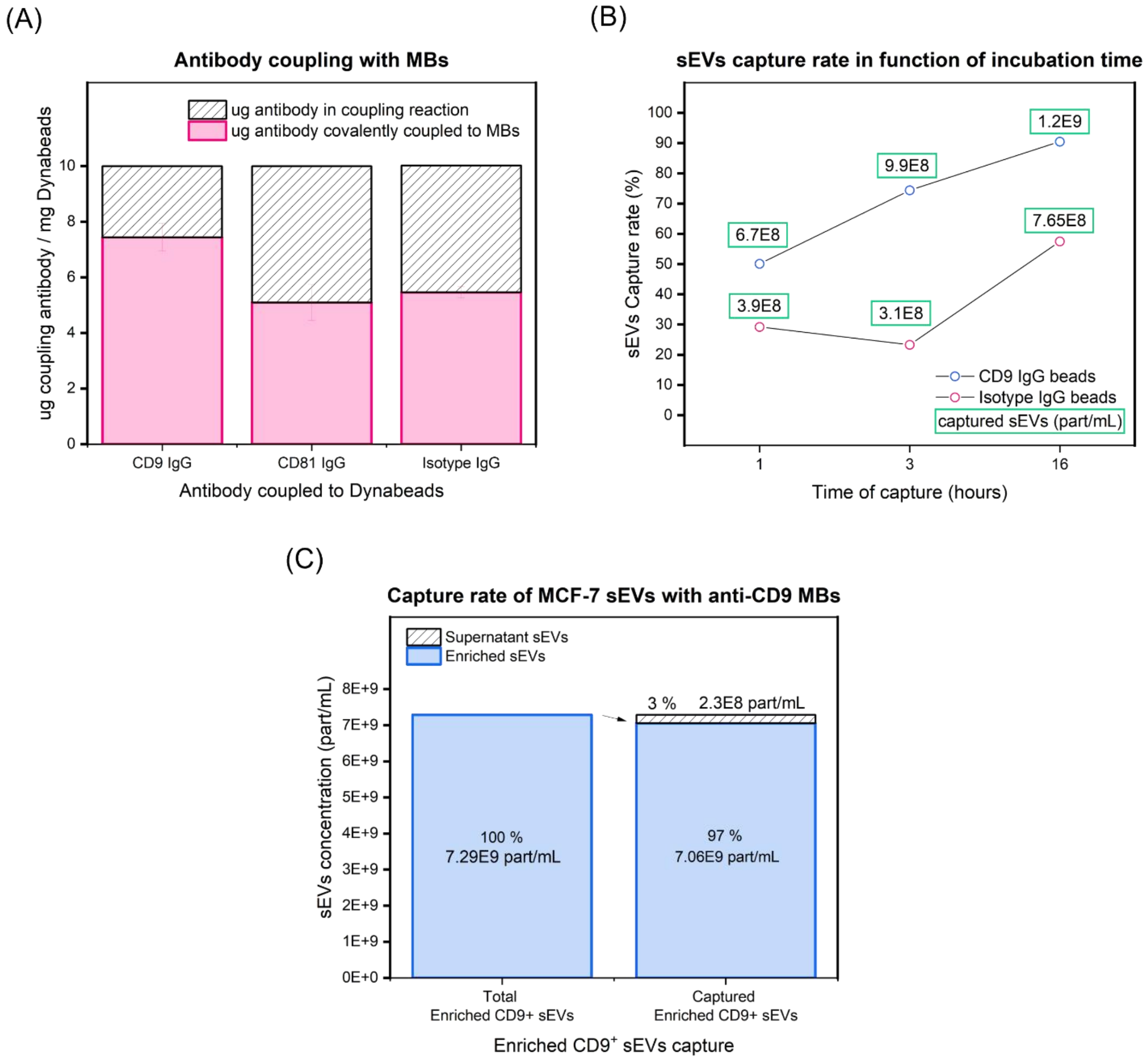

**Fig. S3. Optimization and validation of sEVs capture with Magnetic Beads (MBs).** (A) Determination of antibody coupling efficiency to MBs by the coupling of antibodies anti-CD9, anti-CD81 and isotype IgG. The bars in pink represent the effective amount of antibody coupled to MBs. (B) Determination of MCF-7 sEVs capture efficiency in function of the incubation time by using MBs pre-coated with antibody anti-CD9 and isotype igG. (C) Determination of sEVs capture rate efficiency with MBs anti-CD9. CD9$^{+}$ MCF-7 sEVs were pre-enriched and at a concentration of 7.29E9 part/mL, the sEVs captured in an incubation period of 3 h. The concentration of the remaining sEVs in supernatant was determined by NTA and the capture rate was calculated.

The concentration of eluted CD9$^{+}$ sEVs was determined by NTA and, as before, we captured the sEVs again with MBs anti-CD9 at an initial concentration of 7.29E9 part/mL and determined the concentration of sEVs remaining in the supernatant. We obtained a capture rate of 97 % and concluded that our method shows high efficiency in capturing target sEVs (**Fig. S3-C**). We utilised a quantity of 7.5 µL of the MBs preparations in our assays and, for these conditions, we calculated the corresponding number of MBs and number of sEV capture antibodies per MB (**Table S1**).

If we consider one binding site between the antibodies and sEVs, The Dynabeads in each reaction can capture a theoretical total of 2.2E12 CD9$^{+}$ sEVs or 1.5E12 CD81$^{+}$ sEVs, a quantity significantly larger than what we typically tested.

Also, according to our model, each MB has at least hundred thousand antibodies (4.5E5 anti-CD9 and 3.1E5 anti-CD81 antibodies) supporting our high capture rates.

**Table S1: Determination of number capture antibodies per MB**

| | **MBs anti-CD9** | **MBs anti-CD81** | **MBs isotype IgG** |
|---|---|---|---|
| Volume of MBs per reaction (µL) | 7.5 | 7.5 | 7.5 |
| Mass of MBs per reaction (mg) | 0.075 | 0.075 | 0.075 |
| Number of MBs per reaction | 5.0E+06 | 5.0E+06 | 5.0E+06 |
| Determined mass antibody (µg) per mass MBs (mg) | 7.4 | 5.1 | 5.5 |
| Total antibody mass per reaction (µg) | 0.56 | 0.38 | 0.41 |
| Total antibody moles per reaction (mol) | 3.7E-12 | 2.6E-12 | 2.8E-12 |
| Antibodies molecules per reaction (N) | 2.2E+12 | 1.5E+12 | 1.7E+12 |
| N antibodies per MB | **4.50E+05** | **3.10E+05** | **3.30E+05** |

**Calculations considered:**

- Molecular weight IgG antibody: 150 kDa
- Avogrado constant: 6.022E23 mol$^{-1}$

**4 – Ratio of s-GNPs to sEVs and determination of TEVs and TEPs**

By performing the E-NEXOS assay on the enriched $CD9^{+}CD81^{+}$ sEVs, one can derive a calibration curve, allowing to correlate the concentration of s-GNPs, in an assay, with the concentration of target sEVs (TEVs) (**Fig. 5B**). This is possible as the concentration of TEVs can be established, in these conditions, by techniques like NTA. On the contrary, in mixed sEV samples, i.e. that have not been particularly enriched for a specific population of sEVs, TEVs cannot be determined by NTA and the correlation between s-GNPs and TEVs cannot be established. We used this method of enrichment to determine the correlation between the concentration of s-GNPs and TEVs and applied it to our following experiments.

Additionally, we determined experimentally that no more than 2 s-GNPs could be bound per sEV in our assay configuration (**Fig. 5C**). Furthermore, these results were supported by a theoretical model (**Table S2**).

In the model, we assumed that when a sEV is bound to a MB, 1/4 of the sEV surface area is interacting with the MB, provided by its spherical shape. Additionally, we assumed that per each sEV captured on the surface of the MBs, 1/2 of the sEVs surface could be available for binding to the s-GNPs and that each s-GNP would require at least 1/4 of its surface area to bind to 1/2 of the sEV surface area. With the model, we determined that, in maximum, two s-GNPs could be bound per sEV in our assays.

**Table S2: s-GNPs to sEVs ratio in E-NEXOS**

| | MBs | Mean sEVs | s-GNPs |
|---|---|---|---|
| Diameter (m) | 2.80E-06 | 1.00E-07 | 2.00E-07 |
| Radius (m) | 1.40E-06 | 5.00E-08 | 1.00E-07 |
| Surface Area ($m^2$) | 2.46E-11 | 3.14E-14 | 1.26E-13 |
| 1/4 Surface Area ($m^2$) | N/A | 7.85E-15 | 3.14E-14 |
| 1/2 Surface Area ($m^2$) | | 1.57E-14 | |
| Max number of sEVs bound per MB | | **3136** | |
| Max number of s-GNPs bound per sEV | | | **2** |

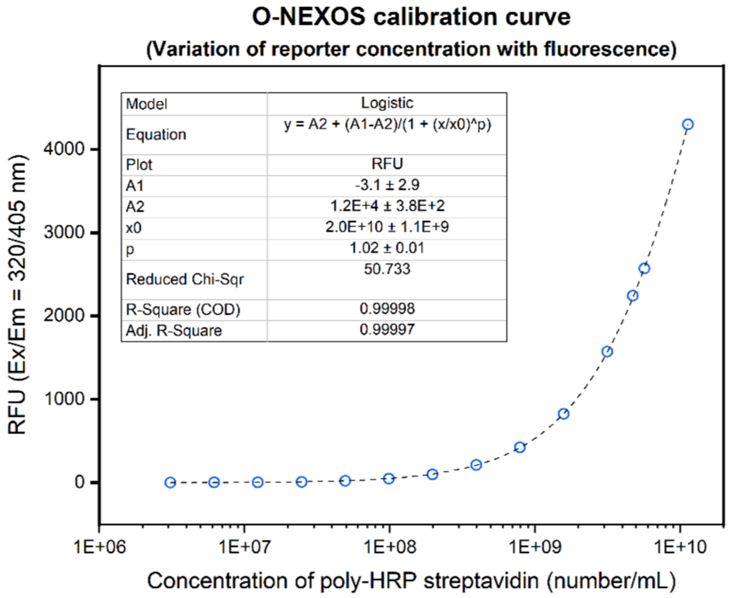

**Fig. S4. Calibration curve of O-NEXOS.** (A) O-NEXOS was calibrated by serially diluting known concentrations of poly-HRP streptavidin and incubating the poly-HRP streptavidin, at the different concentrations, with equal proportions of QuantaBlu™ Peroxidase substrate per each assay. The resulting fluorescence was determined at Ex/Em = 320/405 nm, allowing to correlate the generated fluorescence signal with the concentration of poly-HRP streptavidin per assay (n = 6).

For the determination of target epitopes (TEPs) in O-NEXOS, a calibration curve was derived, allowing to correlate the generated fluorescence signal with the concentration of poly-HRP strep molecules obtained in the reactions (**Fig. S4**).

### 5 – Detection of pre-prepared cancer cell-derived sEVs with NEXOS technology

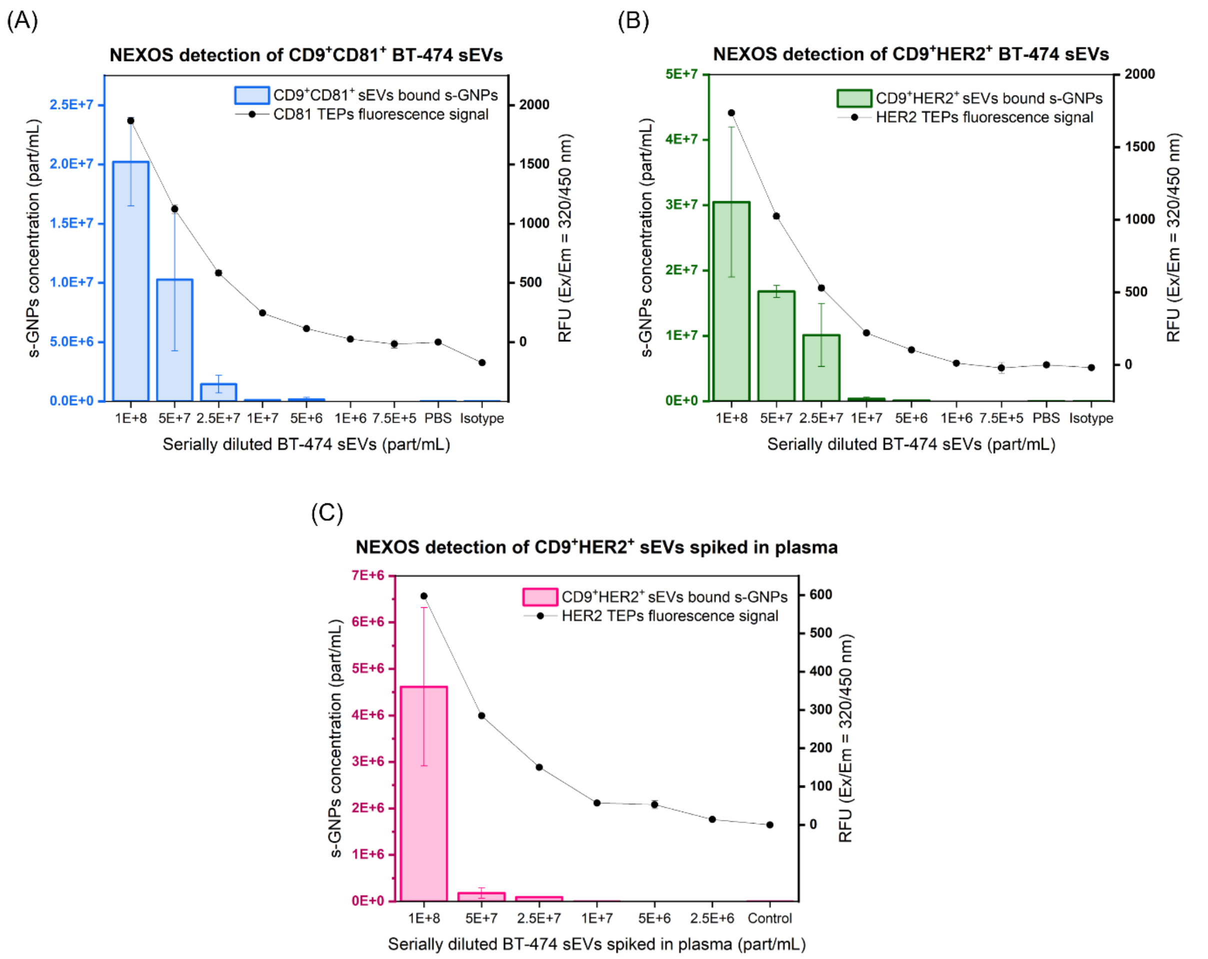

**Fig. S5. Detection of BT-474 sEVs with NEXOS technologies.** Pre-prepared BT-474 sEVs were serially diluted to known concentrations and the following antigens in the conditions described were targeted and detected in E-NEXOS and O-NEXOS: (A) $CD9^+CD81^+$ sEVs in PBS (B) $CD9^+HER2^+$ sEVs in PBS and (C) $CD9^+HER2^+$ sEVs spiked in processed human plasma. Left axes indicate the concentration of s-GNPs bound to sEVs, as determined by E-NEXOS and the right axes indicate the fluorescence obtained, as determined by O-NEXOS (n = 4).

In **Fig. S5**, we show both E-NEXOS and O-NEXOS detection of $CD9^+CD81^+$ BT-474 sEVs in PBS (**Fig. S5-A**), $CD9^+HER2^+$ BT-474 sEVs in PBS (**Fig. S5-B**) and $CD9^+HER2^+$ BT-474 sEVs in processed plasma (**Fig. S5-C**), in units of s-GNPs concentration and RFUs, as complement to the detection of TEVs and TEPs in **Fig. 6**.

**6 – Detection of sEVs with different technologies**

BT-474 sEVs, all from the same batch, were serially diluted to concentrations of 1E10, 5E9, 1E9, 5E8, 1E8, 5E7, 1E7 and 5E6, and compared with the linear detection of NEXOS using BT-474 sEVs. The techniques used were Fluorescence-ELISA, SP-IRIS and IFC as demonstrated in **Fig. S6**.

We determined the calibration curve of ExoELISA-ULTRA CD81 kit using the standards and instructions provided by the supplier (**Fig. S6-A**). Using the serially diluted BT-474 sEVs, ExoELISA-ULTRA CD81 kit only detected sEVs down to a LOD of 5E9 part/mL and a LOQ of 1E10 part/mL since the signal at 5E9 part/mL could not be distinguished from the signal obtained at lower concentrations. Additionally, the signal below 5E9 part/mL could not be distinguished from the blank controls (**Fig. S6-B**).

SP-IRIS detected $CD9^+CD81^+$ and $CD9^+HER2^+$ sEVs at concentrations of 1E9, 1E8 and 1E7 part/mL (**Fig. S6-C** and **Fig. S6-D**). The smallest concentration measured was a preparation of 1E7 part/mL due to the limited chips availability but, upon performing a detection curve with the obtained SP-IRIS measurements, we estimated a LOQ of 9.4E6 part/mL $CD9^+HER2^+$ sEVs and 4.7E6 part/mL of $CD9^+CD81^+$ BT-474, as the minimum concentrations possible to quantify above noise levels originated on the isotype control groups (**Fig. S6-E**).

Finally, IFC detected CD81 and HER2 down to a LOQ of 5E7 part/mL for both markers, albeit detecting more signal overall when targeting for HER2 (**Fig. S6-F**).

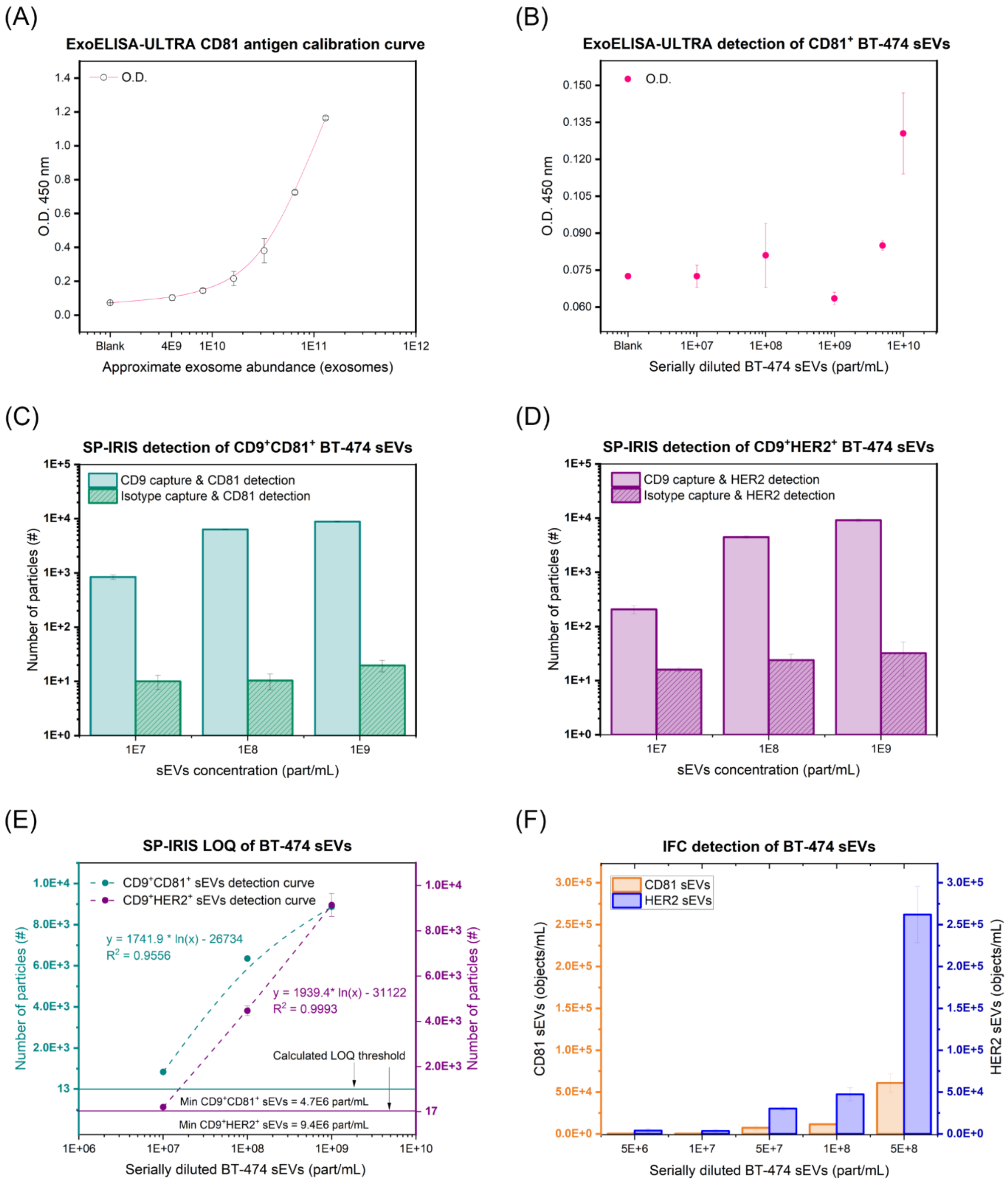

**Fig. S6. Detection of BT-474 sEVs with different technologies.** BT-474 sEVs were pre-prepared and serially diluted to several concentrations as before and the dynamic range and limit in sensitivity of Fluorescence-ELISA, SP-IRIS and IFC was determined. (A) ELISA calibration. (B) CD81 detection on BT-474 sEVs by ELISA. (C) SP-IRIS detection of CD9$^+$CD81$^+$ sEVs. (D) SP-IRIS detection of CD9$^+$HER2$^+$ sEVs. (E) Determination of SP-IRIS LOQ for CD9$^+$CD81$^+$ and CD9$^+$HER2$^+$ sEVs. (F) IFC detection of CD81$^+$ sEVs and HER2$^+$ sEVs.
